# A free boundary mechanobiological model of epithelial tissues

**DOI:** 10.1101/2020.07.02.185686

**Authors:** Tamara A. Tambyah, Ryan J. Murphy, Pascal R. Buenzli, Matthew J. Simpson

## Abstract

In this study, we couple intracellular signalling and cell–based mechanical properties to develop a novel free boundary mechanobiological model of epithelial tissue dynamics. Mechanobiological coupling is introduced at the cell level in a discrete modelling framework, and new reaction–diffusion equations are derived to describe tissue–level outcomes. The free boundary evolves as a result of the underlying biological mechanisms included in the discrete model. To demonstrate the accuracy of the continuum model, we compare numerical solutions of the discrete and continuum models for two different signalling pathways. First, we study the Rac–Rho pathway where cell– and tissue–level mechanics are directly related to intracellular signalling. Second, we study an activator–inhibitor system which gives rise to spatial and temporal patterning related to Turing patterns. In all cases, the continuum model and free boundary condition accurately reflect the cell–level processes included in the discrete model.

## 1 Introduction

Epithelial tissues consist of tightly packed monolayers of cells [1–3]. Mechanical cell properties, such as resistance to deformation and cell size, and chemical cell properties, such as intracellular signalling, impact the shape of epithelial tissues [2, 4]. The role of purely mechanical cell properties on tissue dynamics has been studied using mathematical and computational models [5–10]. Other models focus on intracellular signalling to examine how chemical signalling affects tissue dynamics [11–15]. We extend these studies by developing a model which couples mechanical cell properties to intracellular signalling. We refer to this as *mechanobiological* coupling. By including mechanobiological coupling in a discrete computational framework, new reaction–diffusion equations are derived to describe how cell–level mechanisms relate to tissue–level outcomes.

Epithelial tissues play important roles in cancer development, wound healing and morphogenesis [2,16,17]. Temporal changes in tumour size and wound width in epithelial monolayers can be thought of as the evolution of a free boundary [18, 19]. Many free boundary models use a classical one–phase Stefan condition to describe the evolution of the free boundary [20, 21]. In these models, the speed of the free boundary is proportional to the spatial gradient of the density at the moving boundary [20, 21]. Other free boundary models, particularly those used to study biological development, pre-specify the rate of tissue elongation to match experimental observations [22–28]. In this study, we take a different approach by constructing the continuum limit description of a biologically–motivated discrete model. In doing so, we derive a novel free boundary condition that arises from the underlying biological mechanisms included in the discrete model. While the discrete model is suitable for describing cell–level observations and phenomena [15,29], the continuum limit description is suitable to describe tissue–level dynamics and is more amenable to analysis [30–32].

To confirm the accuracy of the continuum limit description, including the new free boundary condition, we compare the solution of the discrete model with the solution of the continuum model for a homogeneous tissue with no mechanobiological coupling, and observe good correspondence. To investigate mechanobiological coupling within epithelial tissues, the modelling framework is applied in two different case studies. The first case study involves the Rac–Rho pathway where diffusible chemicals called Rho GTPases regulate mechanical cell properties [12, 33–38]. We explicitly consider how the coupling between diffusible chemical signals and mechanical properties lead to different tissue–level outcomes, including oscillatory and non-oscillatory tissue dynamics. The second case study involves the diffusion and reaction of an activator–inhibitor system in the context of Turing patterns on a non-uniformly evolving cellular domain [26–28]. In both case studies, the numerical solution of the continuum model provides an accurate description of the underlying discrete mechanisms.

## 2 Model Description

In this section, we first describe the cell–based model, referred to as the *discrete model*, where mechanical cellular properties are coupled with intracellular signalling. To provide mathematical insight into the discrete model, we then derive the corresponding coarse–grained approximation, which is referred to as the *continuum model*.

### 2.1 Discrete model

To represent a one dimensional (1D) cross section of epithelial tissue, a 1D chain of cells is considered [8,9] (Figure 1). The tissue length, *L*(*t*), evolves in time, while the number of cells, *N*, remains fixed. We define *x*_*i*_(*t*), *i* = 0, 1, … *N*, to represent the cell boundaries, such that the left boundary of cell *i* is *x*_*i*−1_(*t*) and the right boundary of cell *i* is *x*_*i*_(*t*). Epithelial tissues often evolve in confined spaces [39]. Therefore we fix the left tissue boundary at *x*_0_(*t*) = 0, while the right tissue boundary is a free boundary, *x*_*N*_(*t*) = *L*(*t*).

**Figure 1:**
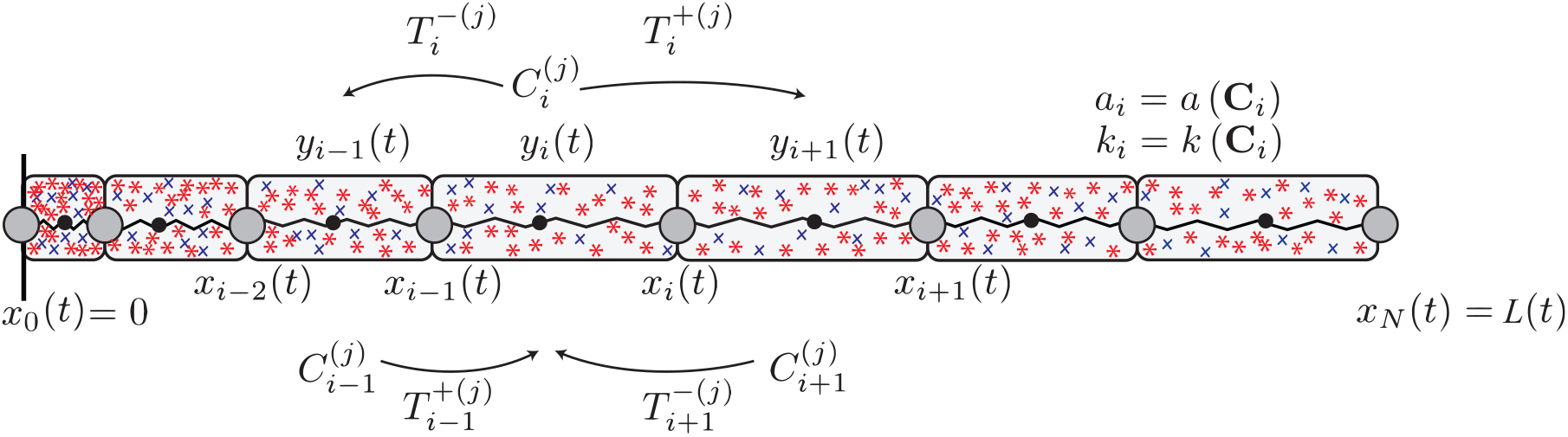
Schematic of the discrete model where mechanical cell properties, *a*_*i*_ and *k*_*i*_, are functions of the family of chemical signals, **C**_*i*_(*t*). In this schematic we consider two diffusing chemical species where the concentration in the *i*^th^ cell is 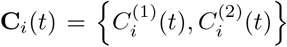. The diffusive flux into cell *i* from cells *i* ± 1, and the diffusive flux out of cell *i* into cells *i* ± 1 is shown. Cell *i*, with boundaries at *x*_*i*−1_(*t*) and *x*_*i*_(*t*), is associated with a resident point, *y*_*i*_(*t*), that determines the diffusive transport rates, 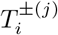.

Each cell, which we consider to be a mechanical spring [8, 9], is assigned potentially distinct mechanical properties, *a*_*i*_ and *k*_*i*_, such that the resulting tissue is heterogeneous (Figure 1) [7]. Each cell *i* contains a family of well–mixed chemical species, 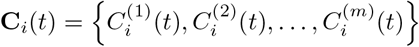, where 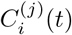 represents the concentration of the *j*^th^ chemical species in cell *i* at time *t*. As the cell boundaries evolve with time, 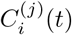 tends to decrease as cell *i* expands. Conversely, 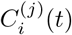 tends to increase as cell *i* compresses. Furthermore, 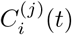 diffuses from cell *i* to cells *i* ± 1. The mechanical properties of individual cells, such as the cell resting length, *a*_*i*_ = *a* (**C**_*i*_), and the cell stiffness, *k*_*i*_ = *k*(**C**_*i*_), may depend on the local chemical concentration, **C**_*i*_(*t*). We refer to this as mechanobiological coupling. The mechanobiological coupling is assumed to be instantaneous as *a*_*i*_ and *k*_*i*_ are functions of the current concentration, **C**_*i*_(*t*).

The motion of each internal cell boundary is governed by Newton’s second law of motion,

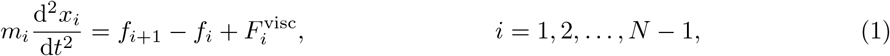

where *m*_*i*_ is the mass associated with cell boundary *i*, 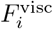 is the viscous force associated with cell boundary *i*, and *f*_*i*_ is the cell-to-cell interaction force acting on cell boundary *i* from the left [5–8]. For simplicity, we choose a linear, Hookean force law given by

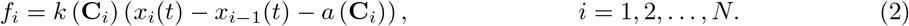

The viscous force is generated by cell–matrix interactions, and is modelled as 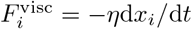 where *η* > 0 is the mobility coefficient [5–8]. As cells move in dissipative environments, the motion of each cell *i* is assumed to be overdamped, such that the left hand side of Equation (1) is zero [6, 7, 40]. Hence, the location of each cell boundary *i* evolves as

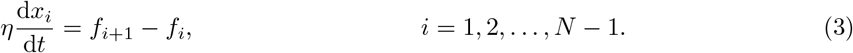

The fixed boundary at *x*_0_(*t*) = 0 has zero velocity, whereas the free boundary at *x*_*N*_(*t*) = *L*(*t*) moves solely due to the force acting from the left:

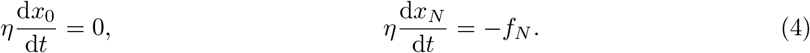

We now formulate a system of ordinary differential equations (ODEs) that describe the rate of change of 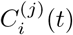 due to changes in cell length and diffusive transport. A position–jump process is used to describe the diffusive transport of 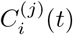. We use 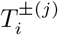 to denote the rate of diffusive transport of 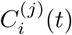 from cell *i* to cells *i* ± 1, respectively [41, 42] (Figure 1). For a standard unbiased position–jump process with a uniform spatial discretisation, linear diffusion at the macroscopic scale is obtained by choosing constant 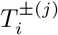 [41]. As the cell boundaries evolve with time, one way to interpret 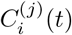 is that it represents a time–dependent, non-uniform spatial discretisation of the concentration profile over the chain of cells. Therefore, care must be taken to specify 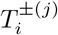 on the temporally evolving spatial discretisation if we suppose the position–jump process corresponds to linear diffusion at the macroscopic level [42].

Yates et al. [42] show that in order for the position–jump process to lead to linear diffusion at the macroscopic level, the length– and time–dependent transport rates must be chosen as

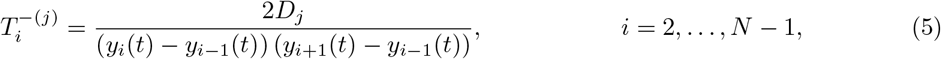

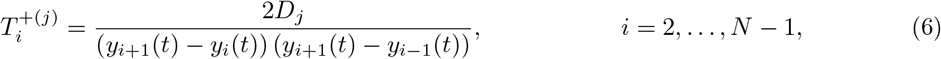

where *D*_*j*_ > 0 is the diffusion coefficient of the *j*^th^ chemical species at the macroscopic level, and *y*_*i*_(*t*) is the *resident point* associated with cell *i* (Figure 1) [42]. The resident points are a Voronoi partition such that the left jump length for the transport of 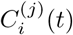 is *y*_*i*_(*t*) − *y*_*i*−1_(*t*), and the right jump length for the transport of 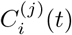 is *y*_*i*+1_(*t*) − *y*_*i*_(*t*) [42]. Complete details of defining a Voronoi partition are outlined in the Appendix B.1.1.

At the tissue boundaries, we set 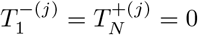 so that the flux of 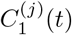 and 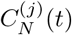 across *x*_0_(*t*) = 0 and *x*_*N*_(*t*) = *L*(*t*) is zero. We follow Yates et al. [42] and choose the inward jump length for the transport of 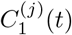 and 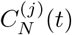 as 2 (*y*_1_(*t*) − *x*_0_(*t*)) and 2(*x*_*N*_(*t*) − *y*_*N*_(*t*)), respectively, giving

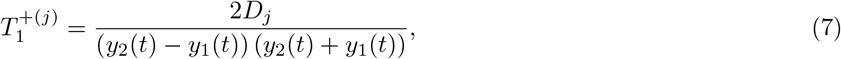

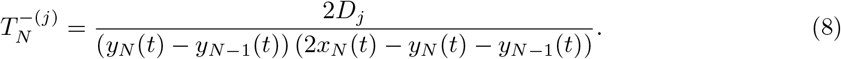

Therefore, the ODEs which describe the evolution of 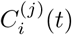 are:

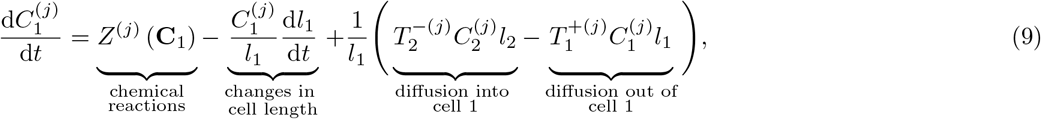

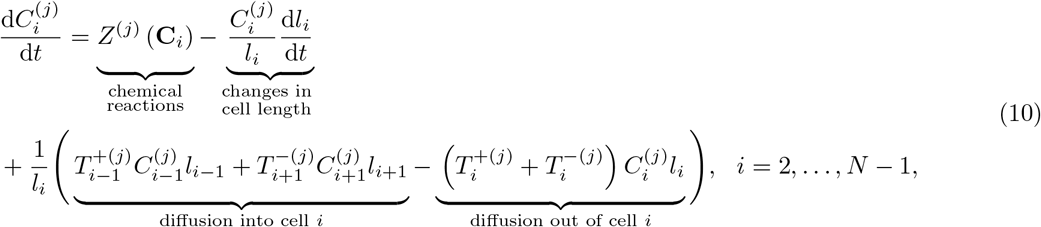

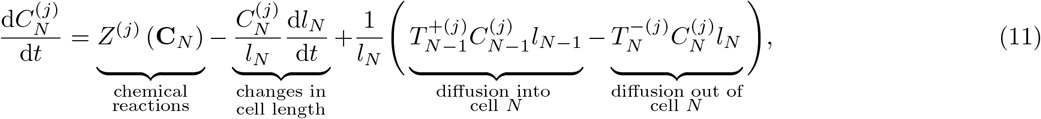

where *l*_*i*_ = *x*_*i*_(*t*) − *x*_*i*−1_(*t*) is the length of cell *i*. Chemical reactions among the chemical species residing in the *i*^th^ cell are described by *Z*^(*j*)^ (**C**_*i*_). The form of *Z*^(*j*)^ (**C**_*i*_) is chosen to correspond to different signalling pathways.

In summary, the discrete model is given by Equations (3)–(11), where Equations (3)–(4) describe the mechanical interaction of cells, and Equations (9)–(11) describe the underlying biological mechanisms. We solve this deterministic system of ODEs numerically using `ode15s` in MATLAB [43]. The numerical method is outlined in the Appendix B.1, and key numerical algorithms to solve the discrete model are available on [GitHub](http://github.com/tamaratambyah/Tambyah2020_v2).

### 2.2 Continuum model

Assuming that the tissue consists of a sufficiently large number of cells, *N*, we now derive an approximate continuum limit description of the discrete model. In the discrete model, *i* = 0, 1, …, *N* is a discrete variable which indexes cell positions and cell properties. The time evolution of the cell boundaries, *x*_*i*_(*t*), is a set of *N* + 1 discrete functions that depend continuously upon time. In contrast, the continuum model describes the spatially continuous evolution of cell boundary trajectories in terms of the cell density per unit length, *q*(*x, t*). In the continuum model, 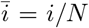 is the continuous analogue of *i* [5]. As *N* → ∞, 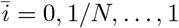 becomes a continuous variable and defines a continuum of cells. The spatially and temporally continuous cell density is [5, 8]

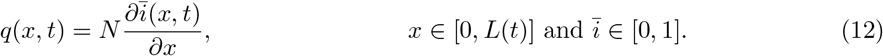

At any time *t*, 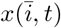 is the inverse function of 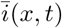 where *x* ∈ [0*, L*(*t*)] for 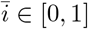. We use 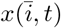 to represent the continuous spatial and temporal evolution of cell boundary trajectories [5].

The discrete quantity **C**_*i*_(*t*) is represented by a multicomponent vector field in the continuum model, 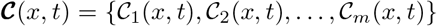. We assume that the mechanical relaxation of cells is sufficiently fast such that the spatial distribution of cell lengths is slowly varying in space [6]. That is, when *k* is sufficiently large, the cells mechanically relax relatively quickly such that each cell in the tissue is approximately the same length. Under this assumption, the location of the resident points can be approximated as the midpoint of each cell,

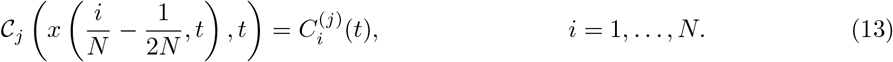

This assumption is further discussed in the Appendix C. In Equation (13), the subscript *j* denotes the *j*^th^ chemical species in the continuum model, and the superscript (*j*) denotes the *j*^th^ chemical species in the discrete analogue. Mechanobiological coupling is introduced by allowing the cell stiffness, 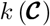, and the cell resting length, 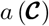, to depend on the local chemical concentration.

We write the linear force law for the continuum of cells as

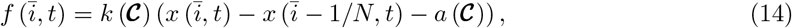

where 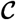 is evaluated at 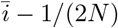. Thus, the equations of motion are:

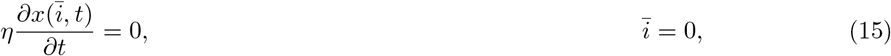

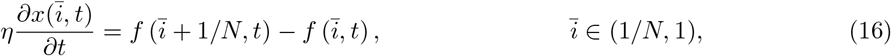

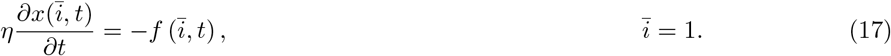

The definition of 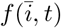 in Equation (14) contains arguments evaluated at 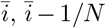 and 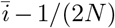. Substituting Equation (14) into Equations (15)–(17), and expanding all terms in a Taylor series about 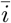 gives,

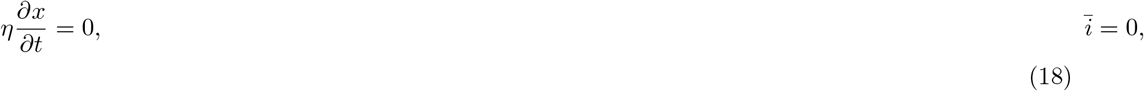

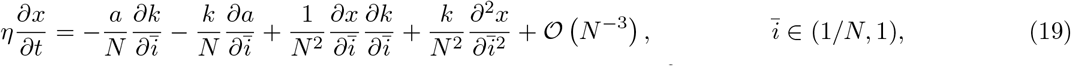

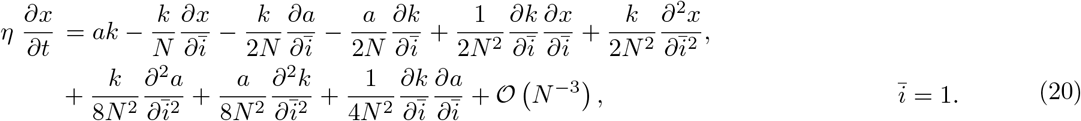

To describe the continuous evolution of cell trajectories and cell properties, we write 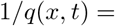 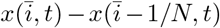 [6, 7], and define the continuous linear force law corresponding to Equation (14) as a 1D stress field,

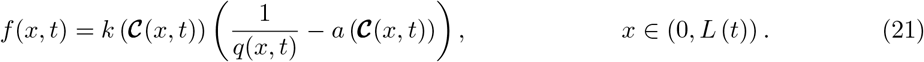

We express Equations (18)–(20) in terms of *q*(*x, t*) and *f* (*x, t*) through a change of variables from 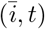 to (*x, t*) [5, 8]. The change of variables gives

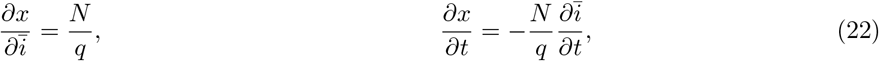

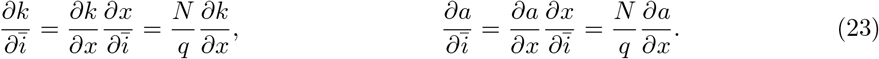

Complete details of the change of variables calculation are outlined in the Appendix A.1.

The local cell velocity, *u*(*x, t*) = *∂x/∂t*, is derived by substituting Equations (22)–(23) into the right hand side of Equation (19). Factorising in terms of *f* (*x, t*) gives,

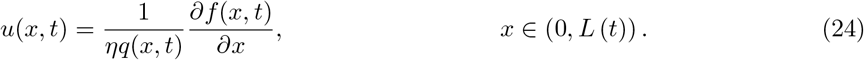

As *u*(*x, t*) = *∂x/∂t*, we substitute Equation (22) into the left hand side of Equation (24) to derive the governing equation for cell density. The resulting equation is differentiated with respect to *x* to give,

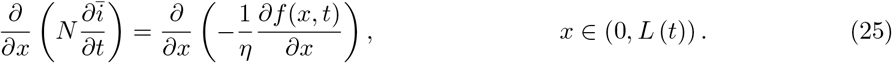

The order of differentiation on the left hand side of Equation (25) is reversed, and Equation (22) is used [5, 8] to give,

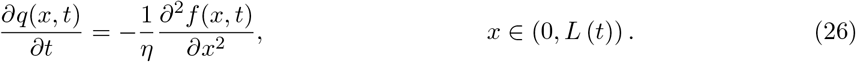

The boundary condition for the evolution of *L*(*t*) is obtained by substituting Equations (22)–(23) into the right hand side of Equation (20), giving

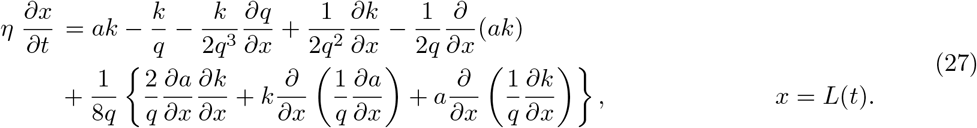

As *u*(*x, t*) = *∂x/∂t*, the left hand side of Equation (27) is equated to Equation (24). Factorising in terms of *f* (*x, t*) gives the free boundary condition

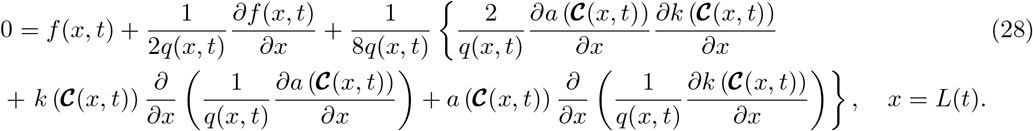

A similar transformation is applied to Equation (18) to yield the left boundary condition as

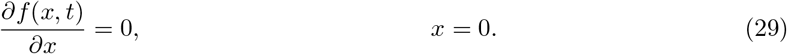

Equations (24), (26), (28) and (29) form a continuum limit approximation of the discrete model. Equations (24), (26) and (29) were reported previously by Murphy et al. [6] who consider heterogeneous tissues of fixed length without mechanobiological coupling. A key contribution here is the derivation of Equation (28), which describes how the free boundary evolves due to the underlying biological mechanisms and heterogeneity included in the discrete model. For a homogeneous tissue where the cell stiffness and cell resting length are constant and independent of 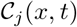, Equation (28) is equivalent to Equation (23) in Baker et al. [5]. The Appendix A.2 shows that Equations (24), (26) and (29) can be derived without expanding all components of Equations (15)–(16). As the definition of 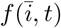 in Equation (14) contains arguments evaluated at 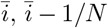 and 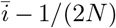, it is necessary to expand all components of Equation (17) about 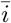 to derive Equation (28). For consistency, Equations (24), (26) and (29) are derived in the same way.

We now consider a reaction–diffusion equation for the evolution of 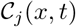. The reaction–diffusion equation involves terms associated with the material derivative, diffusive transport, and source terms that reflect chemical reactions as well as the effects of changes in cell length,

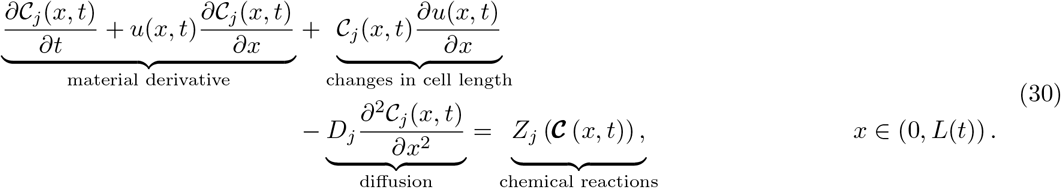

The material derivative arises from differentiating Equation (13) with respect to time, and describes to the propagation of cell properties along cell boundary characteristics [6,7]. The term describing the effects of changes in cell length arises directly from the discrete mechanisms described in Equations (9)–(11). The linear diffusion term arises due to the choice of jump rates in Equations (5)–(8) of the discrete model [42]. Chemical reactions are described by 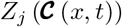, and originate from equivalent terms in the discrete model, *Z*^(*j*)^ (**C**_*i*_).

Boundary conditions for 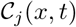 are chosen to ensure mass is conserved at *x* = 0 and *x* = *L*(*t*). As the left tissue boundary is fixed, we set 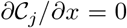 at *x* = 0. At *x* = *L*(*t*), we enforce that the total flux of 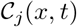 in the frame of reference co-moving with the right tissue boundary is zero for all time,

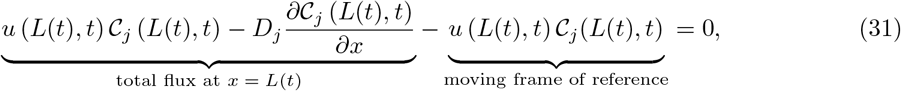

where *u* (*L*(*t*)*, t*) = d*L/*d*t*. Thus, 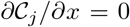 at *x* = *L*(*t*). Writing Equation (30) in conservative form gives,

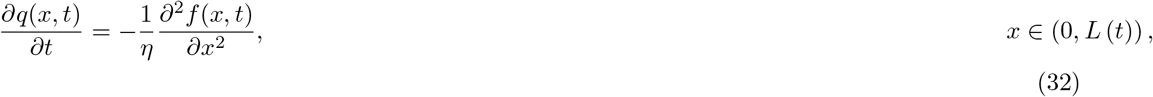

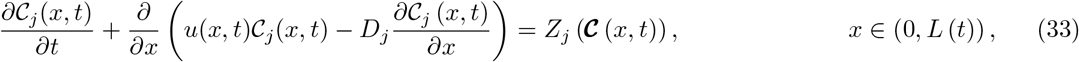

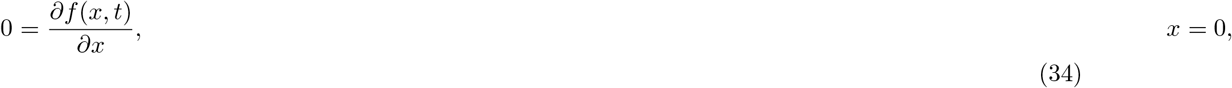

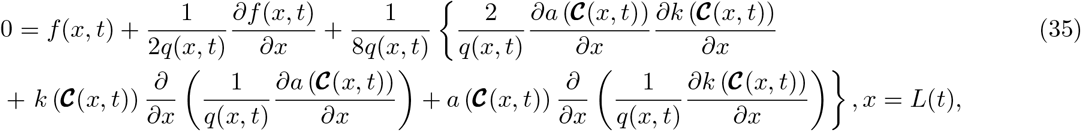

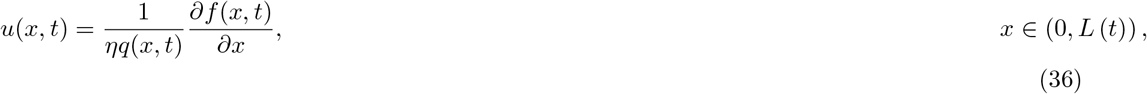

where

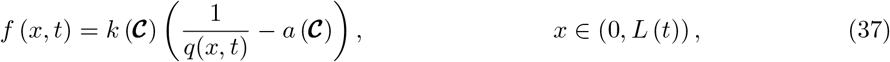

and 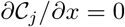 at *x* = 0 and *x* = *L*(*t*).

Equations (32)–(37) are solved numerically using a boundary fixing transformation [44]. In doing so, Equations (32)–(37) are transformed from an evolving domain, *x* ∈ [0*, L*(*t*)], to a fixed domain, *ξ* ∈ [0, 1], by setting *ξ* = *x/L*(*t*) [44]. The transformed equations are discretised using a standard implicit finite difference method with initial conditions *q*(*x,* 0) and 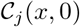. The numerical method is outlined in the Appendix B.2, and key numerical algorithms to solve the continuum model are available on [GitHub](http://github.com/tamaratambyah/Tambyah2020_v2).

## 3 Results

To examine the accuracy of the new free boundary model, we compare solutions from the discrete and continuum models for epithelial tissues consisting of *m* = 1 and *m* = 2 chemical species.

### 3.1 Preliminary results: Homogeneous tissue

In all simulations, an epithelial tissue with just *N* = 20 cells is considered. Figure 2 in Baker et al. [5] shows that the accuracy of the continuum model increases with *N*. We present similar results in the Appendix C, for *N* > 20, to confirm this. Thus, choosing a relatively small value of *N* is a challenging scenario for the continuum model. In the discrete model, each cell *i* is initially the same length, *l*_*i*_(0) = 0.5, such that *L*(0) = 10. The discrete cell density is *q*_*i*_(*t*) = 1/*l*_*i*_(*t*) which corresponds to *q*(*x,* 0) = 2 for *x* ∈ [0*, L*(*t*)] in the continuum model.

**Figure 2:**
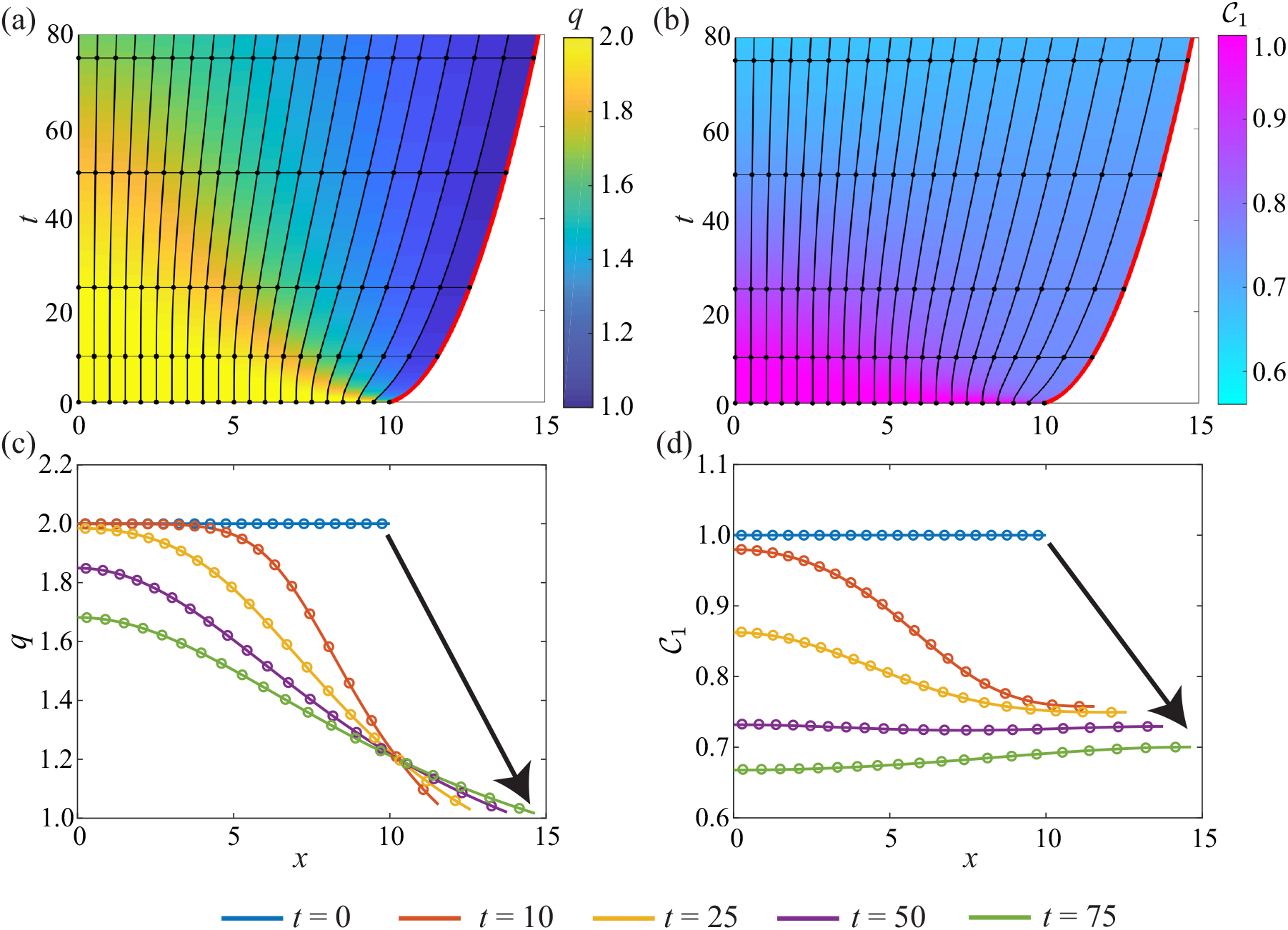
Homogeneous tissue with *N* = 20 cells and one chemical species where 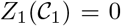, and *a* = *k* = *D*_1_ = *η* = 1. Characteristic diagrams in (a)–(b) illustrate the position of cell boundaries where the free boundary is highlighted in red. The colour in (a)–(b) represents *q*(*x, t*) and 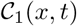 respectively. In (a)–(b), the black horizontal lines indicate times at which *q*(*x, t*) and 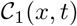 snapshots are shown in (c)–(d). In (c)–(d), the discrete and continuum solutions are compared as the dots and solid line respectively for *t* = 0, 10, 25, 50, 75 where the arrow represents the direction of time.

The simplest application of the free boundary model is to consider cell populations without mechanobiological coupling. For a homogeneous tissue with one chemical species 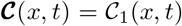, the cell stiffness and cell resting length are constant and independent of 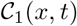. Thus, the governing equations for *q*(*x, t*) and 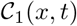 are only coupled through the cell velocity, *u*(*x, t*). To investigate how non-uniform tissue evolution affects the chemical concentration of cells, we set 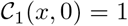 for *x* ∈ [0*, L*(*t*)] and 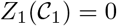.

Figure 2(a) demonstrates a rapid decrease in the cell density at *x* = *L*(*t*) as the tissue relaxes and the cells elongate. This decreases the chemical concentration (Figure 2(b)). As the tissue mechanically relaxes, the cell boundaries form a non-uniform spatial discretisation on which 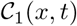 is transported (Figure 2(a)–(b)). Figure 2(c)–(d) demonstrates that the continuum model accurately reflects the biological mechanisms included in the discrete model, despite the use of truncated Taylor series expansions. Additional results in the Appendix D show that *q*(*x, t*), 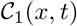 and *L*(*t*) become constant as *t* → ∞.

### 3.2 Case study 1: Rac–Rho pathway

We now apply the mechanobiological model to investigate the Rac–Rho pathway. Rho GTPases are a family of signalling molecules that consist of two key members, RhoA and Rac1. Rho GTPases cycle between an active and inactive state, and regulate cell size and cell motility [12, 35, 37, 45, 46]. Additionally, Rho GTPases play roles in wound healing [47] and cancer development [48, 49]. New experimental methods [50, 51] have discovered a connection between cellular mechanical tension and Rho GTPase activity [52]. Previous studies focus on using a discrete modelling framework to investigate this relationship, and conclude that epithelial tissue dynamics is dictated by the strength of the mechanobiological coupling [12, 33]. Experimental evidence suggests that some GTPases can move between cells [53]. We extend previous mathematical models which consider mechanobiological coupling [12,33] by incorporating diffusive chemical transport and chemical variation associated with changes in cell length.

To investigate the impact of mechanobiological coupling on epithelial tissue dynamics, we let 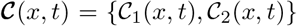 such that 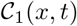 is the concentration of RhoA and 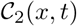 is the concentration of Rac1. In the discrete and continuum models, cells are assumed to behave like linear springs [8, 9]. Thus, cellular mechanical tension is defined as the difference between the length and resting length of cells. Mechanobiological coupling is proportional to cellular tension, and is included in 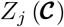. As Rho GTPase activity increases cell stiffness [54], we choose 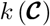 as an increasing function of either 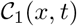 or 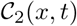. Furthermore, 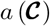 is chosen to reflect the fact that RhoA promotes cell contraction [12, 33]. We first set *D*_1_ = *D*_2_ = 1, and then vary the diffusivity to explore how this affects the tissue–level dynamics.

The effect of RhoA is considered first, where 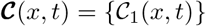. We include the same mechanobiological coupling as [12, 33] and let,

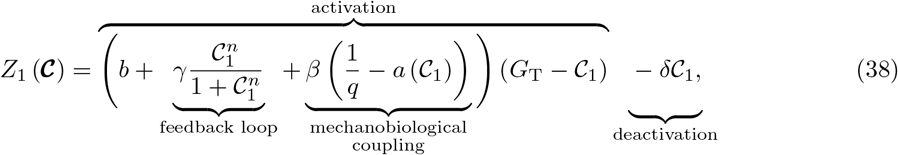

where *b* is the basal activation rate, *G*_T_ is the total amount of active RhoA, and *δ* is the deactivation rate [12, 33]. The activation term describes a positive feedback loop, governed by *γ*, to reflect the fact that RhoA self–activates [12, 33]. Mechanobiological coupling, governed by *β*, is proportional to mechanical tension.

Similar to [12], we find that the tissue either mechanically relaxes or continuously oscillates depending on the choice of *β* (Figure 3). Figure 3(a),(c) illustrates non-oscillatory tissues for weak mechanobiological coupling, *β* = 0.2, whereas increasing the strength of the mechanobiological,*β* = 0.3, leads to temporal oscillations in the tissue length and sharp transitions between high and low levels of RhoA (Figure 3(b),(d)). Figure 3(e)–(h) illustrates that the continuum model accurately describe non-oscillatory and oscillatory tissue dynamics.

**Figure 3:**
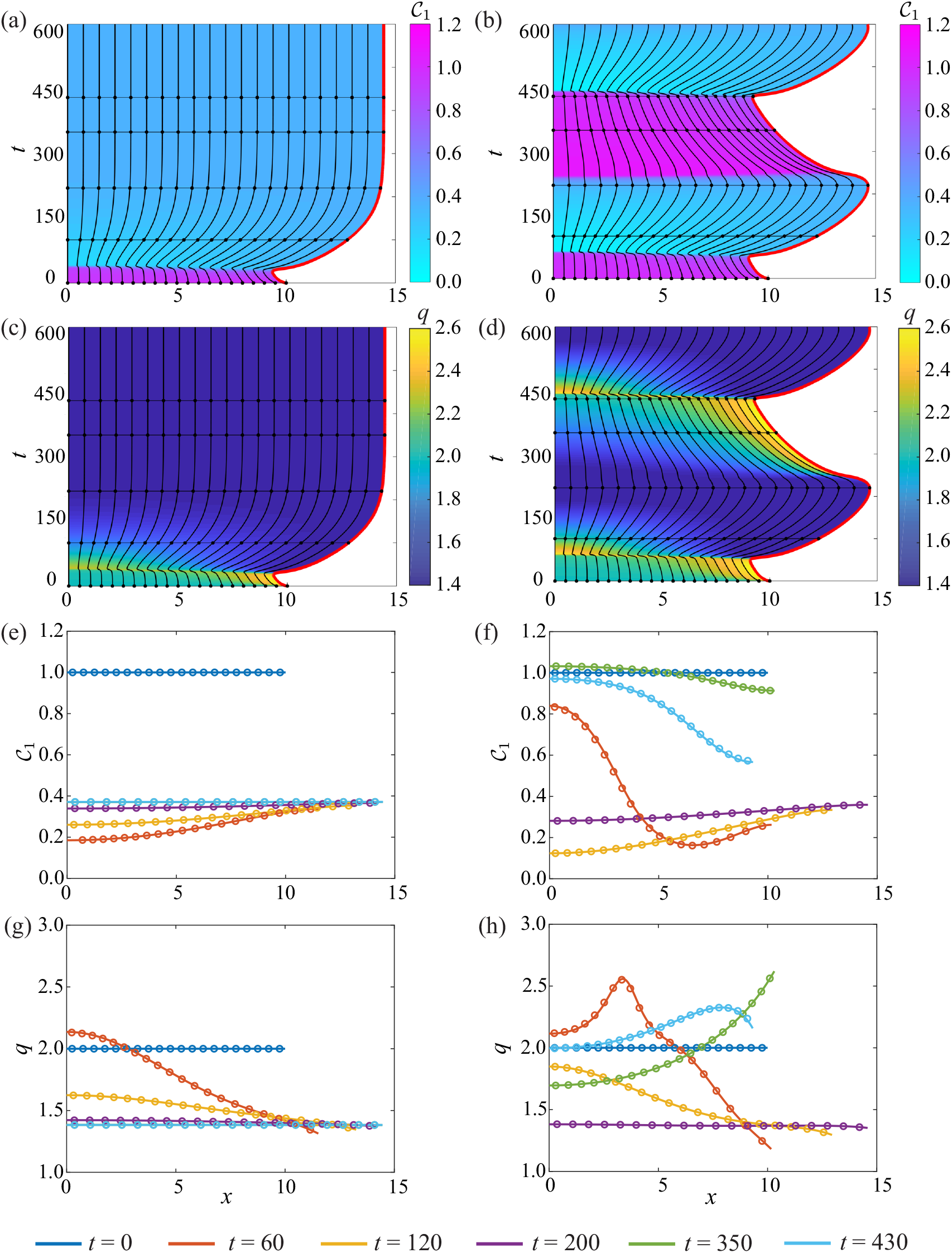
1D tissue dynamics where RhoA is coupled to mechanical cell tension. (a),(c),(e),(g) correspond to *β* = 0.2 and (b),(d),(f),(h) relate to *β* = 0.3. Characteristic diagrams in (a)–(d) illustrate the evolution of cell boundaries where the free boundary is highlighted in red. The colour in (a)–(b) represents 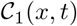 and *q*(*x, t*) in (c)–(d). The black horizontal lines indicate times at which 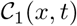 and *q*(*x, t*) snapshots are shown in (e)–(f) and (g)–(h) respectively. In (e)–(h), the discrete and continuum solutions are compared as the dots and solid line respectively for *t* = 0, 100, 220, 350, 430. In both systems, 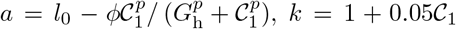, *D*_1_ = 1, *η* = 1, and 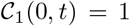 for *x* ∈ [0*, L*(*t*)]. Parameters: *b* = 0.2, *γ* = 1.5, *n* = 4, *p* = 4, *G*_T_ = 2, *l*_0_ = 1, *ϕ* = 0.65, *G*_h_ = 0.4, *δ* = 1.

Diffusion is often considered to be stabilising [55] so we hypothesise that increasing *D*_1_ could smooth the tissue oscillations. Surprisingly, Figure 4 shows that as *D*_1_ increases we observe a phase shift but no change in the amplitude of oscillations. This test case provides further evidence of the ability of the new continuum model to accurately capture key biological mechanisms included in the discrete model, even when *D*_1_ is relatively small (Figure 4). As with any numerical solution of a reaction–diffusion equation, a finer computational mesh is required to solve the continuum model when the diffusivity is small [56], meaning that the continuum model becomes computationally expensive as the diffusivity decreases. Under these conditions we prefer to use the discrete model.

**Figure 4:**
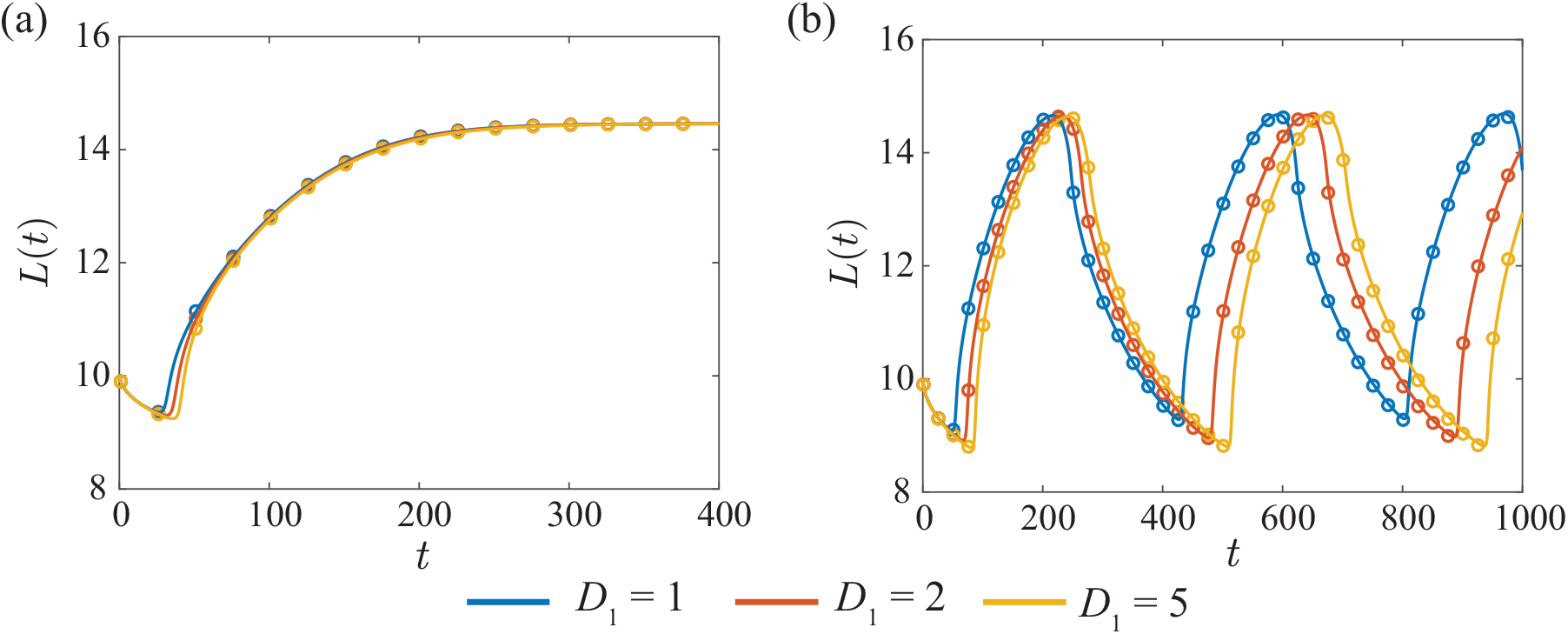
The effect of diffusion on the dynamics of the free boundary for (a) a non-oscillatory system with *β* = 0.2 and (b) a oscillatory system with *β* = 0.3. The discrete solution is shown as the dots and the continuum solution as solid line. Parameters are as in Figure 3.

To examine the combined effect of RhoA and Rac1 on epithelial tissue dynamics, we let 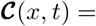 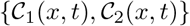 and consider [12],

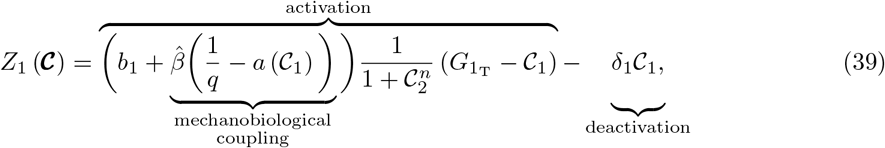

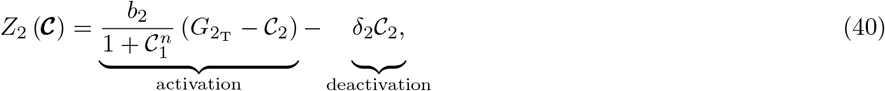

where 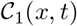 is the concentration of RhoA and 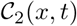 is the concentration of Rac1. In a weakly coupled system when 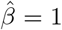, we observe the fast transition from the initial concentrations of RhoA and Rac1 to the steady state concentration (Figure 5(a),(c)). Analogous to Figure 3(a),(c), temporal oscillations arise when the mechanobiological coupling is strong, 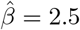 (Figure 5(b),(d)).

**Figure 5:**
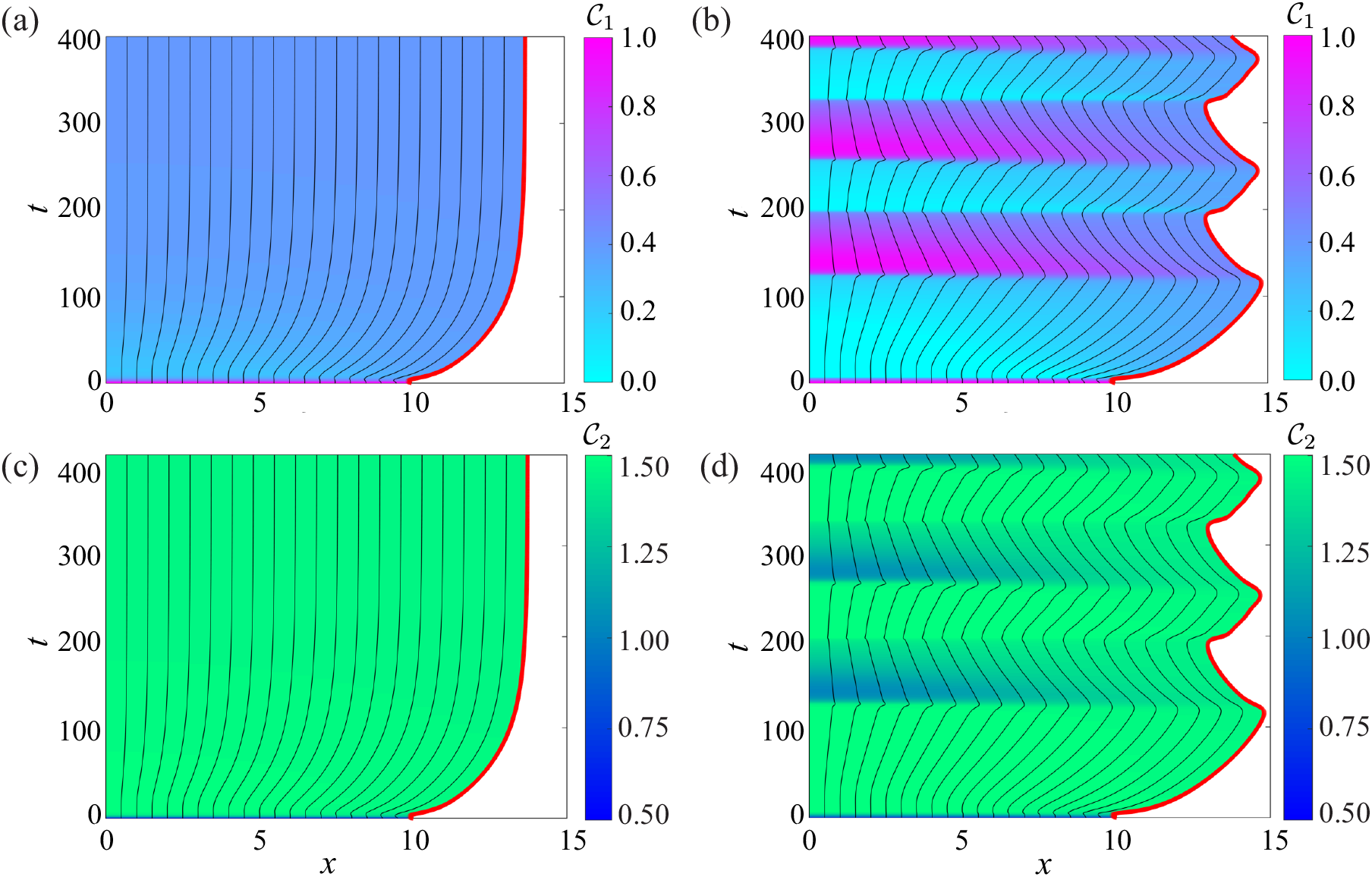
Characteristic diagrams for the interaction of RhoA and Rac1 where the free boundary is highlighted in red. (a),(c) correspond to a non-oscillatory system where 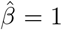 and (b),(d) relate to an oscillatory system where 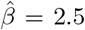. The colour in (a)–(b) denotes the concentration of RhoA and the concentration of Rac1 in (c)–(d). In both systems, 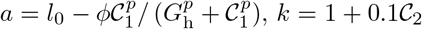, *D*_1_ = *D*_2_ = 1 and *η* = 1. The initial conditions are 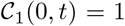 and 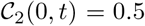 for *x* ∈ [0*, L*(*t*)]. Parameters: *b*_1_ = *b*_2_ = 1, *δ*_1_ = *δ*_2_ = 1, *n* = 3, *p* = 4, 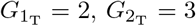, *l*_0_ = 1, *ϕ* = 0.65, *G*_h_ = 0.4. Discrete and continuum solutions are compared in the Appendix E.

The mechanobiological coupling and intracellular signalling in Figures 3–5 extend upon previous Rho GTPase models [12, 33]. We note that the source terms in Equations (38)–(40) can be applied to a single, spatially–uniform cell to explore how the stability of the equilibria depend on model parameters [12]. This analysis, outlined in the Appendix E.1, was used to inform our choices of *β* and 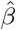.

### 3.3 Case study 2: Activator–inhibitor patterning

Case study 2 considers an activator–inhibitor system [55, 57, 58]. Previous studies of activator–inhibitor patterns on uniformly growing domains have characterised the pattern splitting and frequency doubling phenomena which occur naturally on the skin of angelfish [25–28, 59]. To investigate how diffusion driven instabilities arise on a non-uniformly evolving domain, we let 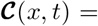 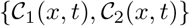 with *D*_1_ ≠ *D*_2_, and use Schnakenberg kinetics,

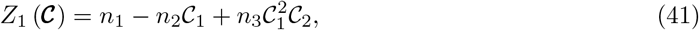

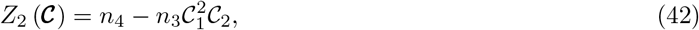

where 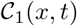 is the activator and 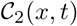 is the inhibitor [55]. The parameters, *n*_*i*_ > 0 for *i* = 1, 2, 3, 4, govern activator–inhibitor interactions. Non-dimensionalisation of Equations (26) and (30) reveals that the linear stability analysis is analogous to the classical stability analysis of Turing patterns on fixed domains [55]. Thus, we define the relative diffusivity, *d* = *D*_2_/*D*_1_, with the expectation that there exists a critical value, *d*_*c*_, that depends upon the choice of *n*_*i*_, such that diffusion driven instabilities arise for *d > d*_*c*_ [55, 57]. The linear stability analysis and computation of *d*_*c*_ is outlined in the Appendix F.1.

A homogeneous tissue is initialised to investigate the affect of *d* on the evolution of activator–inhibitor patterns. As the mechanical cell properties are chosen to be constant and independent of 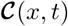, the governing equations for *q*(*x, t*), 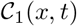 and 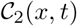 are only coupled through the cell velocity, *u*(*x, t*). Thus, the tissue evolves non-uniformly as a result of this coupling. For *d < d*_*c*_, the distribution of 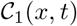 and 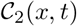 varies in time but remains approximately spatially uniform throughout the tissue (Figure 6(a),(c),(e),(g)). Thus, only temporal patterning arises when *d < d*_*c*_. Figure 6(b),(d),(f),(h) demonstrates that spatial–temporal patterns develop when *d > d*_*c*_. Similar to [25–27], we observe splitting in activator peaks for *d > d*_*c*_ where the concentration of 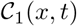 is at a minimum. Figure 6(b) shows that two distinct activator peaks arise. The long time behaviour of the tissue is examined in the Appendix F. Figure 6(e),(g) shows excellent agreement between the solutions of the discrete and continuum models when *d < d*_*c*_, whereas Figure 6(f),(h) shows a small discrepancy between the solutions of the discrete and continuum models when *d > d*_*c*_.

**Figure 6:**
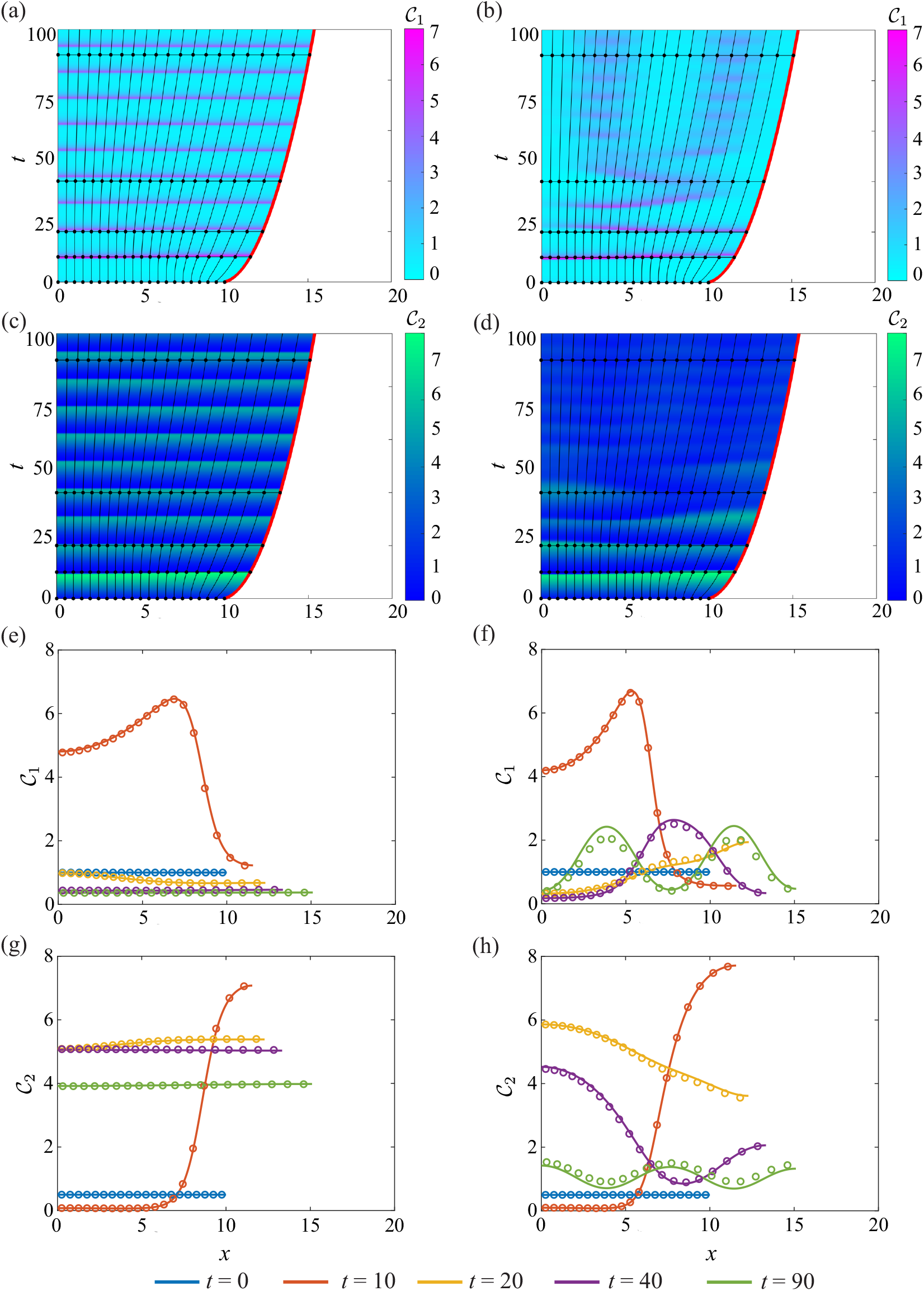
Activator–inhibitor patterns in a homogeneous tissue. In (a),(c),(e),(g), *D*_1_ = 2 and *D*_2_ = 3 with *d < d*_*c*_. In (b),(d),(f),(h), *D*_1_ = 0.5 and *D*_2_ = 5 with *d > d*_*c*_. (a)–(d) shows the cell boundaries where the free boundary is highlighted in red. The colour in (a)–(b) represents 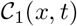 and 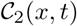 in (c)–(d). The black horizontal lines indicate times at which 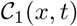 and 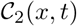 snapshots are shown in (e)–(f) and (g)–(h) respectively. In (e)–(h), the discrete (dots) and continuum (solid line) solutions are compared at *t* = 0, 10, 20, 40, 90, 150. In both systems, 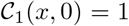 and 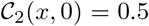 for *x* ∈ [0*, L*(*t*)] and *a* = *k* = *η* = 1 Parameters: *n*_1_ = 0.1*, n*_2_ = 1*, n*_3_ = 0.5*, n*_4_ = 1 and *d*_*c*_ = 4.9842

As the mechanical relaxation in Figure 6(f),(h) is relatively fast, *k* = 1, the small discrepancy between the solutions of the discrete and continuum models decreases as the tissue mechanically relaxes. Additional results in Figure 7 confirm that when the mechanical relaxation is slower, *k* = 0.5, the discrepancy between solutions of the discrete and continuum models is more pronounced, as expected. Thus, the continuum model accurately describes tissue–level behaviour when the mechanical relaxation of cells is sufficiently fast. In cases where the mechanical relaxation is slow, we advise use of the discrete model.

**Figure 7:**
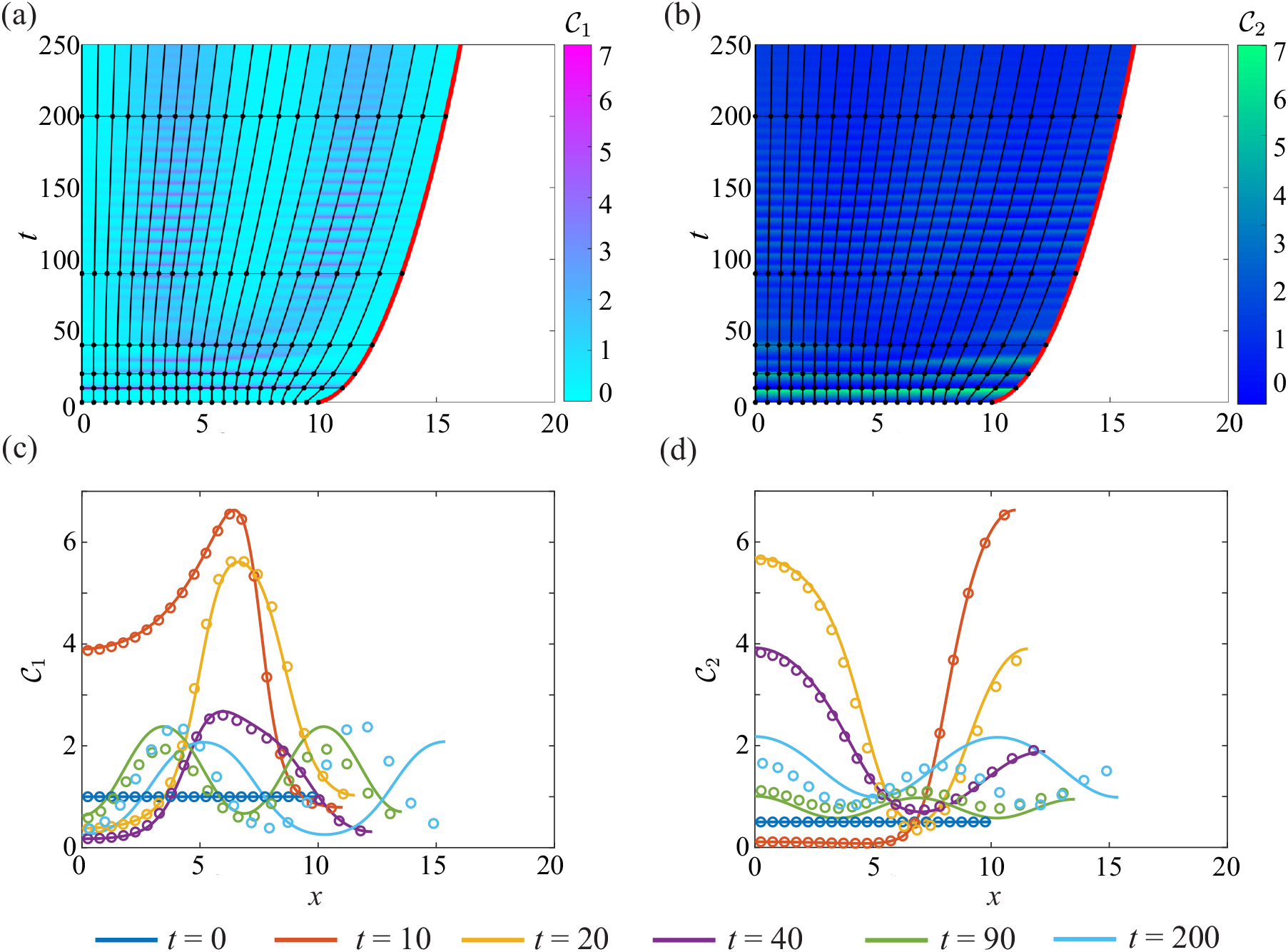
The evolution of spatial–temporal patterns in a homogeneous tissue with Schnakenberg dynamics, where *d > d*_*c*_ and *k* = 0.5. Characteristic diagrams in (a)–(b) illustrate the evolution of cell boundaries where the free boundary is highlighted in red. The colour in (a) represents 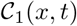 and 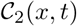 in (b). The black horizontal lines indicate times at which 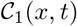 and 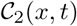 snapshots are shown in (c) and (d) respectively. In (c)–(d), the discrete and continuum solutions are compared as the dots and solid line respectively for *t* = 0, 10, 20, 40, 90, 200. The initial conditions are 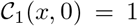 and 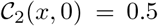 for *x* ∈ [0*, L*(*t*)], with *a* = *η* = 1, *D*_1_ = 0.5 and *D*_2_ = 5. Parameters: *n*_1_ = 0.1*, n*_2_ = 1*, n*_3_ = 0.5*, n*_4_ = 1 and *d*_*c*_ = 4.9842.

## 4 Conclusion

In this study, we present a novel free boundary mechanobiological model to describe epithelial tissue dynamics. A discrete modelling framework is used to include mechanobiological coupling at the cell level. Tissue–level outcomes are described by a system of coupled, non-linear partial differential equations, where the evolution of the free boundary is governed by a novel boundary condition. The evolution of the free boundary is an emergent property rather than being pre-specified to match experimental observations [22–28]. By constructing the continuum limit of the biologically–motivated discrete model, we arrive at a novel free boundary condition which describes how mechanobiological coupling dictates epithelial tissue dynamics. In deriving the continuum model, we make reasonable assumptions that *N* is sufficiently large, and that the time scale of mechanical relaxation is sufficiently fast [7]. Case studies involving a homogeneous cell population, the Rac–Rho pathway and activator–inhibitor patterning demonstrate that the continuum model reflects the biologically–motivated discrete model even when *N* is relatively small. These case studies show how non-uniform tissue dynamics, including oscillatory and non-oscillatory tissue behaviour, arises due to mechanobiological coupling.

There are many potential avenues to generalise and extend the modelling framework presented in this study. We take the most fundamental approach and use a linear force law to describe mechanical relaxation. One avenue for extension would be to explore non-linear force laws [5, 9]. Another choice we make is to suppose that chemical transport is described by linear diffusion, whereas other choices, such as drift-diffusion, are also possible [60]. We also implement the most straightforward assumption that the mechanobiological coupling is instantaneous, and a further extension would be to include delays in the mechanobiological coupling. Incorporating delays could be useful to represent chemical processes which occur on different time scales to the mechanical relaxation [14, 61]. We always consider the case where the evolution of the free boundary is driven by mechanical relaxation. However, it is also possible to consider a more general scenario where the evolution of the position of cell boundaries explicitly depends on the local chemical concentration [11, 26]. Further extensions of the free boundary model would be to introduce cell proliferation and cell death [5, 7, 62]. These extensions are relatively straightforward to incorporate into the discrete model, and the tissue–level dynamics could be analysed by constructing a new continuum model [5, 7].

The free boundary model presented in this study is tractable because we consider a 1D cross section of epithelial tissue. This simplification is relevant when we consider slender tissues [64]. A significant extension of our work would be to consider a full two dimensional (2D) discrete model by representing cells as polygons [11, 15, 29]. To derive a 2D continuum model, we would need to invoke several simplifying assumptions about the cell-to-cell interactions and connectivity [1, 65]. A 2D free boundary continuum model would require more sophisticated numerical methods to solve as the boundary fixing transformation used in this study would no longer be appropriate. Therefore, we leave the question of developing a 2D model for future consideration.

## Data access

MATLAB implementations of key numerical algorithms is available on [GitHub](http://github.com/tamaratambyah/Tambyah2020_v2).

## Author Contributions

All authors contributed equally to the design of the study. TAT performed numerical simulations with the assistance of RJM. TAT drafted the article and all authors gave approval for publication.

## Acknowledgements

This work is supported by the Australian Research Council (DP200100177, DP180101797). We acknowledge the computational resources provided by the High Performance Computing and Research Support Group. We thank the two anonymous referees for their helpful comments.

## Appendices

## A Continuum model derivation

## A.1 Change of variables

This section outlines the change of variables from 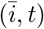 to (*x, t*) used in Section 2.2. Rewriting Equation (12), the cell density per unit length, *q*(*x, t*), is

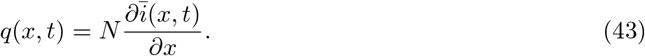

To perform a change of variables from 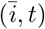 to (*x, t*), we calculate the Jacobian of the coordinate transformation [5, 8, 9],

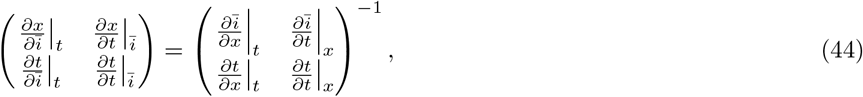

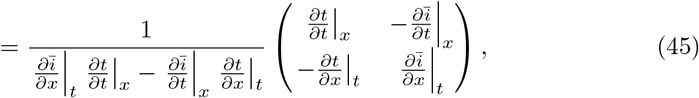

to arrive at the relationships,

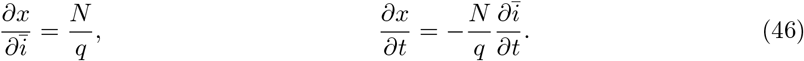

Using the chain rule, the second derivatives are,

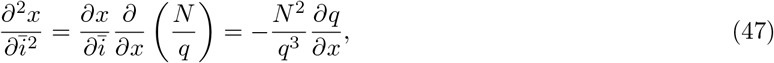

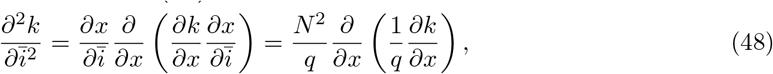

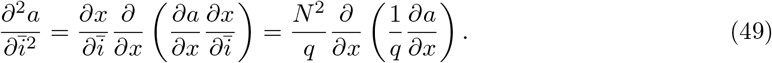

## A.2 Derivation of governing equation for cell density in factorised form

Here we outline how Equations (24), (26) and (29) can be obtained using the factorised form of 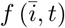 in Equation (14). The equations of motion in Equations (15)–(17) are restated as,

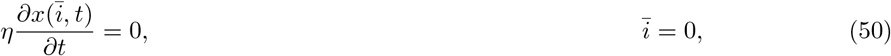

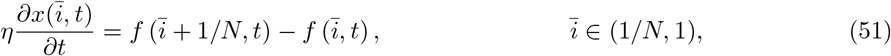

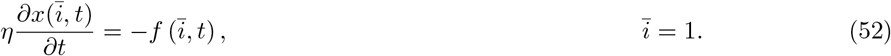

To derive the local cell velocity in Equation (24), the right hand side of Equation (51) is expanded in a Taylor series about 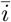,

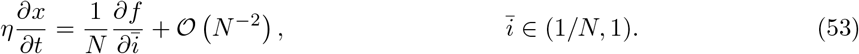

Neglecting non-zero higher order terms and using the chain rule gives,

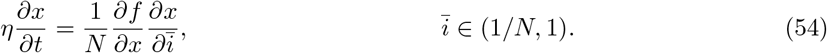

Equation (46) is substituted into the right hand side of Equation (54) to derive the local cell velocity, *u*(*x, t*) = *∂x/∂t*, as,

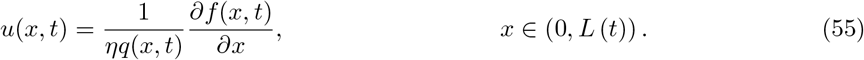

As *u*(*x, t*) = *∂x/∂t*, Equation (46) is substituted into the left hand side of Equation (55). Differentiating the resulting equation with respect to *x* gives,

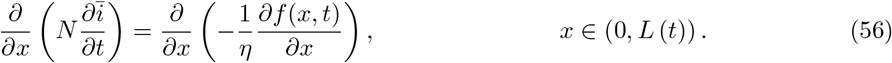

The order of differentiation on the left hand side of Equation (56) is reversed, and Equation (46) is used to give,

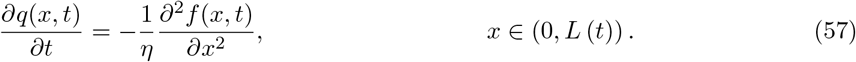

To derive the left boundary condition in Equation (29), the left hand side of Equation (50) is equated to Equation (55) giving,

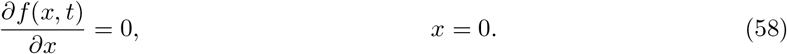

Thus, we have shown that Equations (24), (26) and (29) can be obtained using the factorised form of 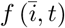 in Equation (14), and first order Taylor series expansions. In the main document we do not pursue this approach. This is because more care is required to obtain the correct form of the free boundary equation in Equation (28). Extensive numerical exploration confirms that the approach taken in the main document is necessary to reflect the underlying biological mechanisms included in the discrete model.

## B Numerical methods

This section outlines the numerical method used to solve the discrete and continuum models with one chemical species, *m* = 1. Key numerical algorithms for *m* = 1 and *m* = 2 chemical species are available on [GitHub](http://github.com/tamaratambyah/Tambyah2020_v2).

## B.1 Discrete model

For *N* cells and one chemical species, the discrete model consists of 2*N* + 1 ordinary differential equations. For simplicity, we write **C**= *C*^(1)^ = *C* and *D*_1_ = *D*. Equations (3)–(11) are restated as:

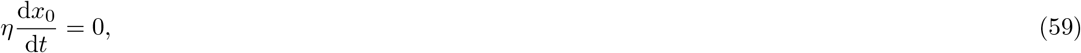

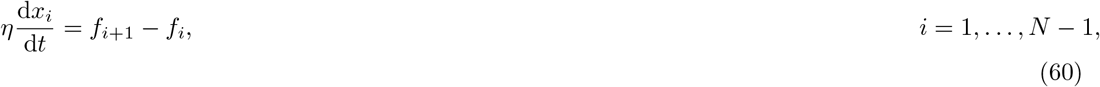

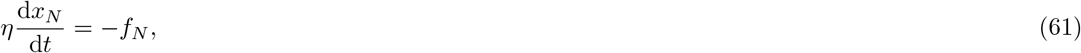

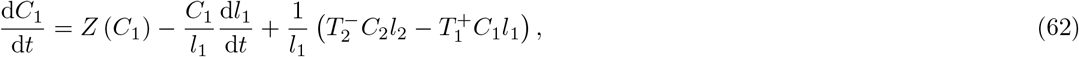

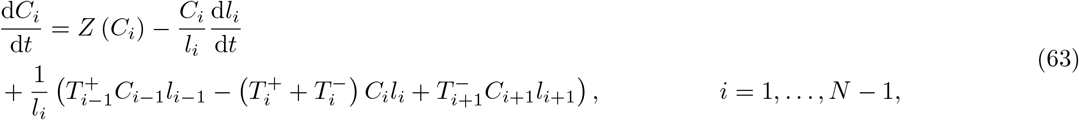

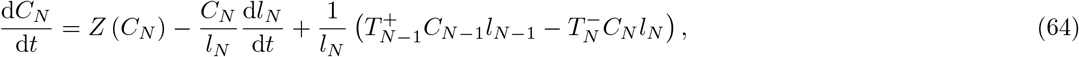

where *l*_*i*_ = *x*_*i*_(*t*) − *x*_*i*−1_(*t*) is the length of cell *i* and

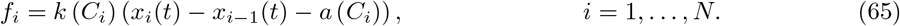

The transport rates for internal cells are

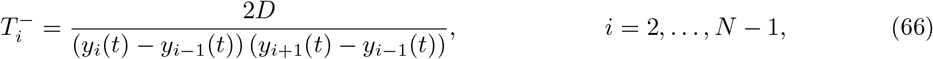

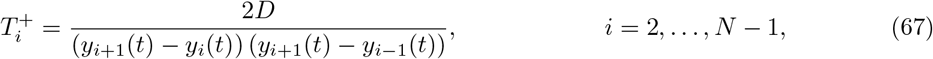

and

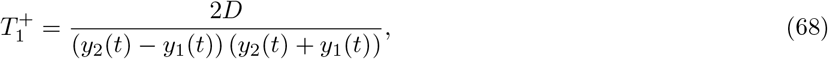

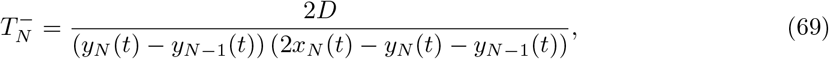

and 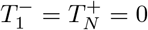 for boundary cells [42]. Equations (59)–(69) are solved numerically using `ode15s` in MATLAB [43]. At each time step, we use a Voronoi partition to compute the resident points, *y*_*i*_(*t*), and the transport rates, 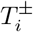.

## B.1.1 Voronoi partition

To define a Voronoi partition, we set the resident point of cell 1, *y*_1_(*t*), as the midpoint of its respective cell boundaries,

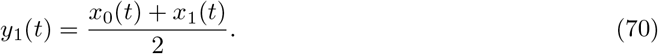

The Voronoi partition enforces that the cell boundaries correspond to the midpoints of resident points [42]. Thus, the following relationship holds

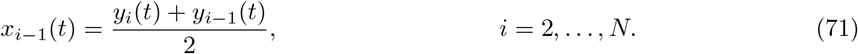

Equations (70) and (71) can be written as the following system of linear equations,

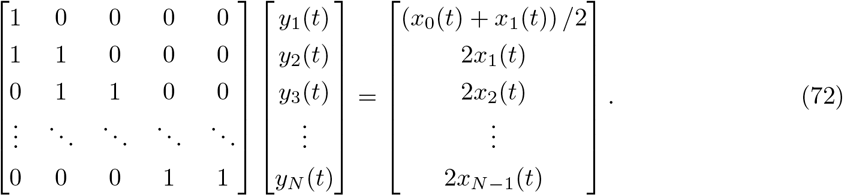

Equation (72) is solved numerically for the resident points at each time step of the discrete simulation. As Equation (72) is a lower triangular matrix system, we use the Thomas Algorithm [56].

## B.2 Continuum model

We now outline the numerical method used to solve the continuum model and write 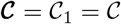 and *D*_1_ = *D* for simplicity. Equations (32)–(37) are restated as a system of coupled, non-linear partial differential equations:

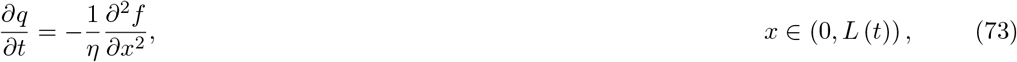

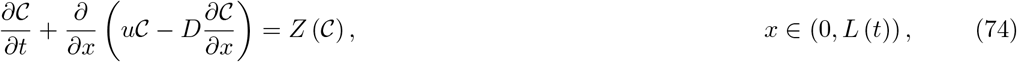

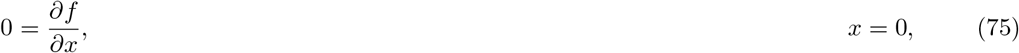

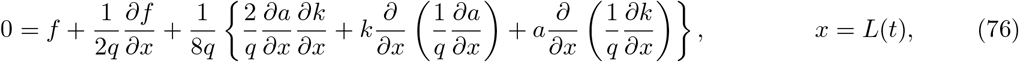

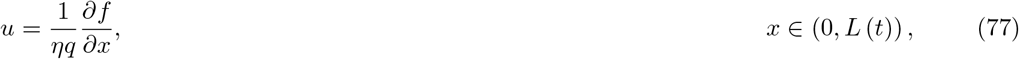

where

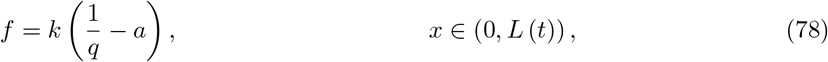

and 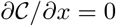 at *x* = 0 and *x* = *L*(*t*).

A standard boundary fixing transformation is used to transform Equations (73)–(78) from an evolving domain, *x* ∈ [0*, L*(*t*)], to a fixed domain, *ξ* ∈ [0, 1], by setting *ξ* = *x/L*(*t*) [44]. Invoking this transform yields:

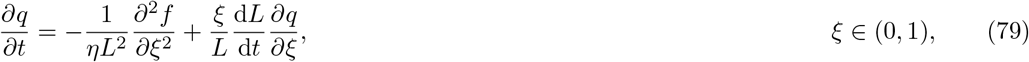

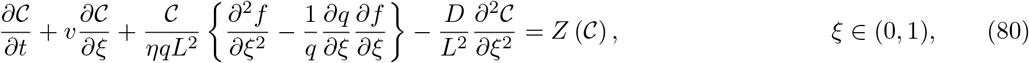

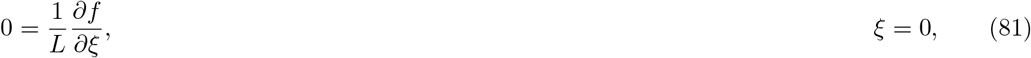

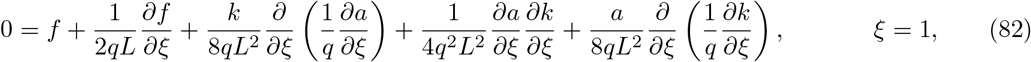

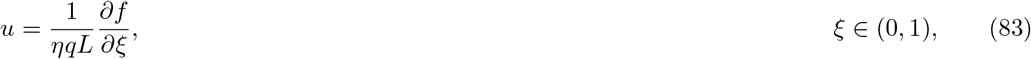

where

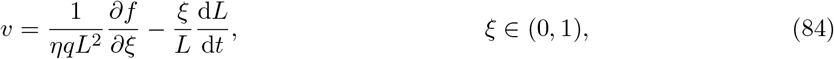

and 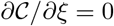 at *ξ* = 0 and *ξ* = 1.

Equations (79)–(84) are spatially discretised on a uniform mesh, with *n* = 1/Δ*ξ* + 1 nodes. The value of *q*(*ξ, t*) and 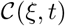 at the *i*^th^ spatial node and the *j*^th^ temporal node are approximated as 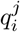 and 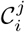 respectively, where *ξ* = (*i* − 1)Δ*ξ* and *t* = *j*Δ*t*. A standard implicit finite difference method is used to approximate spatial and temporal derivatives [56].

We first consider the discretisation of the governing equation and boundary conditions for cell density. Central difference approximations are used to discretise Equation (79) as

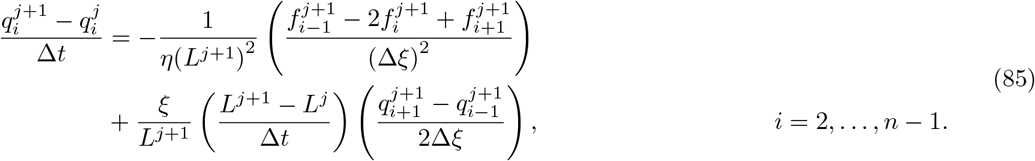

Equations (81) and (82) are discretised using appropriate forward and backward difference approximations,

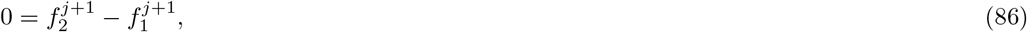

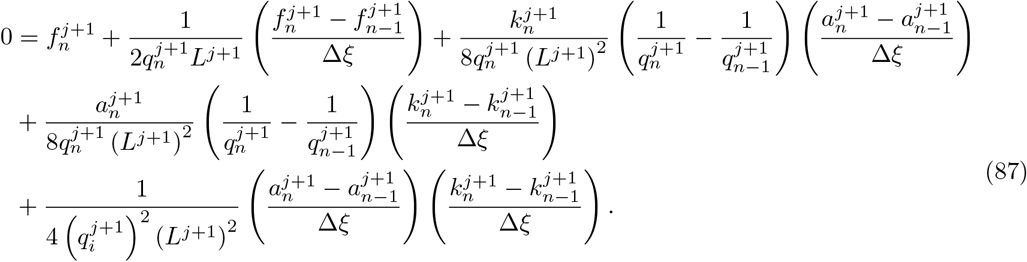

We now consider the discretisation of the governing equation and boundary conditions for 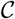. As 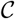 propagates along cell boundary characteristics, Equation (84) is used to upwind the first derivative of 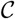 in Equation (80). For *v*_*i*_ *>* 0, a backward difference approximation is used to approximate the first derivative of 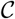, and all other spatial derivates are approximated using central difference approximations,

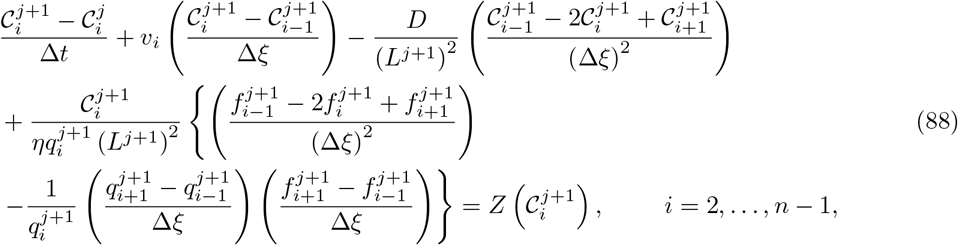

where

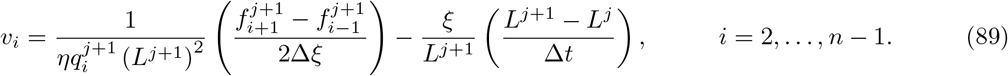

Similarly, forward difference approximations are used when *v*_*i*_ < 0. The boundary condition at *ξ* = 0 is

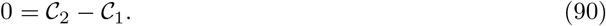

Numerical exploration revealed that a ghost node was necessary to solve 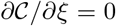 at *ξ* = 1. The use of a ghost node ensured that the numerical solution of the continuum model agreed with the solution of the discrete model. The ghost node is placed outside the right domain boundary at *i* = *n* + 1. A central difference approximation is applied to the zero–flux boundary condition to obtain 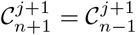. To incorporate the ghost node, Equation (80) is factorised as

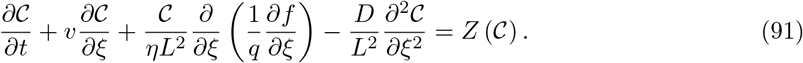

Backward and central difference approximations are used to discretise Equation (91) as

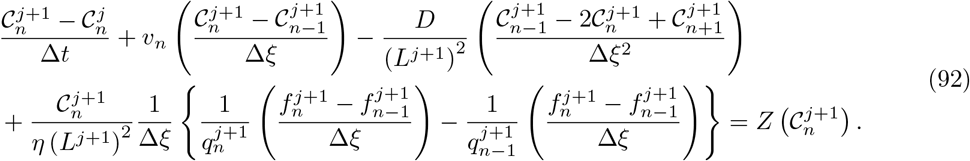

Substituting 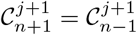 into Equation (92) and factorising gives

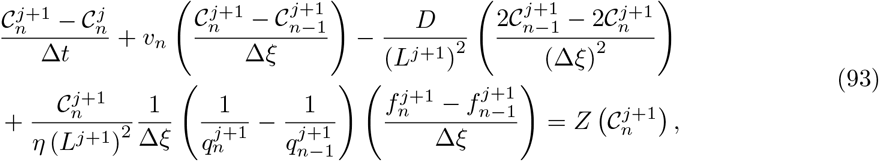

where

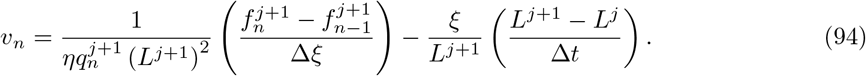

Equation (78) is discretised as

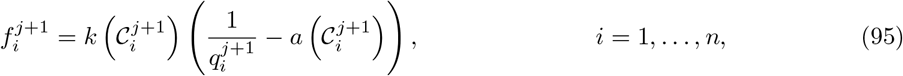

and substituted into Equations (85)–(94) to form a non-linear system of equations.

We solve Equations (85)–(87) for 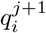 and Equations (88)–(90),(93)–(94) for 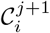 using the Newton-Raphson method [66]. At each Newton-Raphson iteration, Equation (83) is used to update the position of the free boundary as

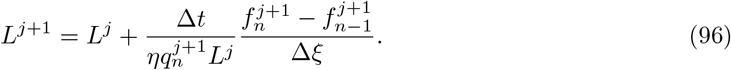

Newton-Raphson iterations are continued until the norm of the difference between successive solution estimates of 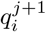 and 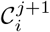 are both less than a specified tolerance, *ϵ*. To ensure all numerical results are grid–independent, Δ*ξ* = 10^*−*3^, Δ*t* = 10^*−*3^ and *ϵ* = 10^*−*8^. A finer computational mesh is required when *D <* 1. All linear systems are solved using the Thomas algorithm [56].

## C Accuracy of the continuum model

In deriving the continuum model, we assume that the tissue consists of a sufficiently large number of cells, and that the mechanical relaxation is sufficiently fast. This section examines the consequences of these assumptions by investigating the effect of *N* and *k* on the accuracy of the continuum model. The infinity norm is chosen as the error metric to compare solutions of the discrete and continuum models. The error metric is used to compare the free boundary position as predicted by the discrete model, *L*_dis_(*t*), and the continuum model, *L*_ctm_(*t*).

In all simulations, we consider a homogeneous tissue with one chemical species, 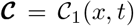, and no chemical source, 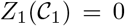. The initial conditions are *q*(*x,* 0) = 2 and 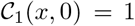 for *x* ∈ [0*, L*(0)]. The effect of *N* on the accuracy of the continuum model is determined by setting *k* = 1, and choosing *L*(0) = *N/*2 for *N* = 5, 20, 40, 60, 75 such that *q*(*x,* 0) = 2. To determine the effect of *k* on the accuracy of the continuum model, we set *N* = 20 with *L*(0) = 10, and choose *k* = 0.5, 1, 1.5, 2. As varying *k* changes the time scale in which the tissue mechanically relaxes, all simulations are conducted to time *t* where *L*(*t*) = 1.5*L*(0).

Figure 8 illustrates that the error metric decreases as *N* and *k* become sufficiently large. Thus, the continuum model is not valid and may not accurately reflect the discrete model when the number of cells is small, and the mechanical relaxation is slow. Under these parameter regimes, we advise use of the discrete model.

**Figure 8:**
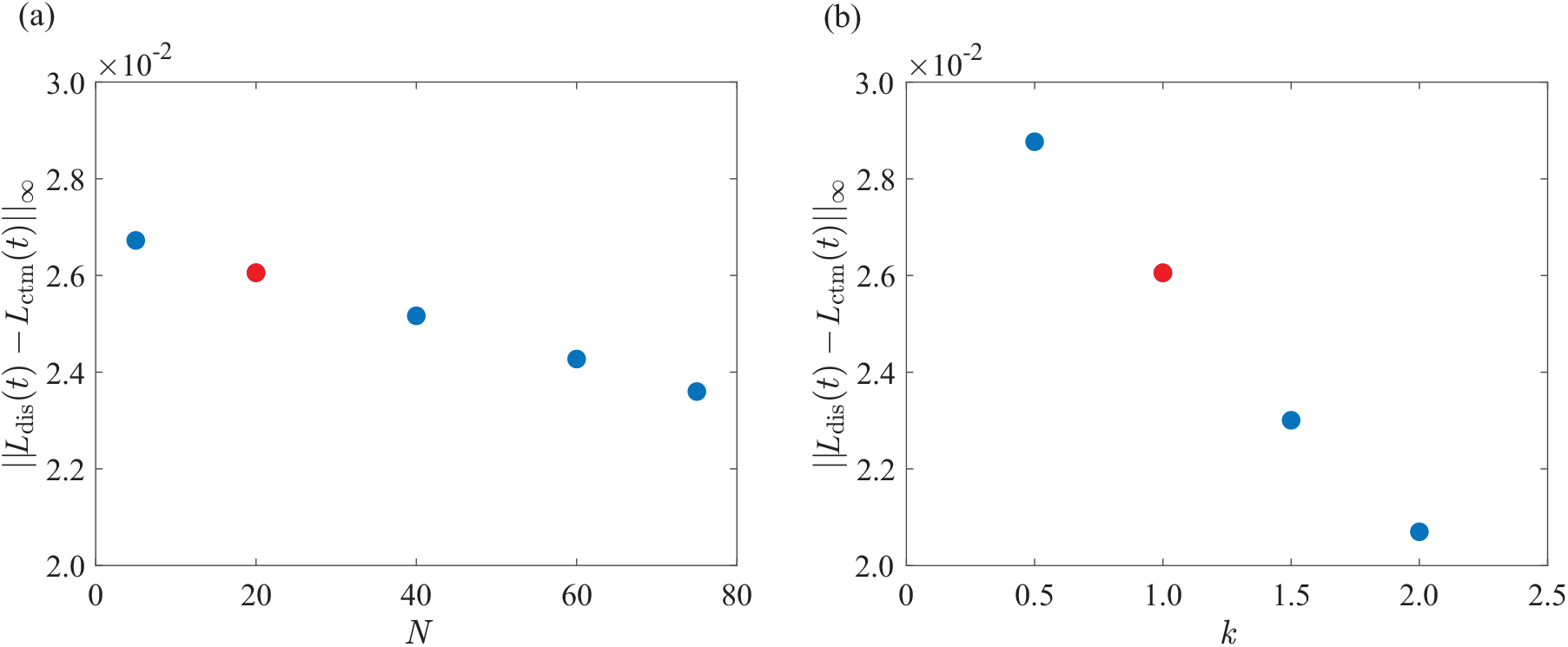
Comparison of the error metric for the free boundary position for a homogeneous tissue with *a* = *D*_1_ = *η* = 1. In (a), *N* is varied and *k* = 1 remains constant. In (b), *k* is varied and *N* = 20 remains constant. The red dot corresponds to the error metric for *N* = 20 and *k* = 1.

## D Preliminary results: Homogeneous tissue

In Section 3.1, we examine a homogeneous tissue with *N* = 20 cells, one chemical species, 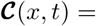 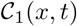, and no chemical source, 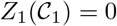. Figure 9(a)–(b) demonstrates that *q*(*x, t*), 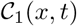 and *L*(*t*) become constant as *t* → ∞. To obtain expressions for the long time behaviour of *q*(*x, t*), 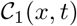 and *L*(*t*), the total number of cells, *N*, and total number of chemical particles, *P*, are computed as,

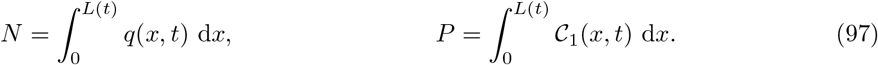

**Figure 9:**
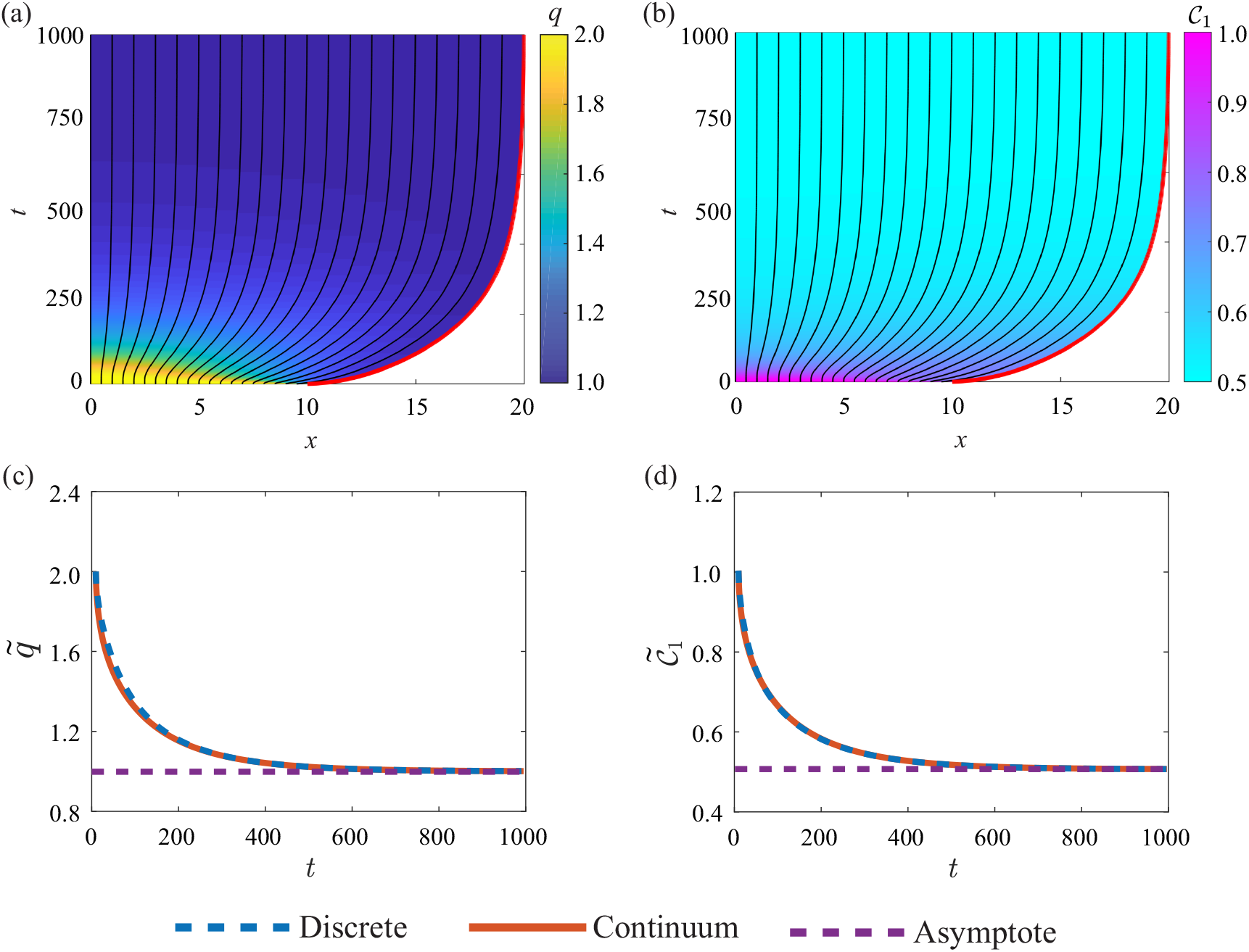
Homogeneous tissue consisting of one chemical species where 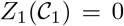 and *a* = *k* = *D*_1_ = *η* = 1. Characteristic diagrams in (a)–(b) illustrate the position of cell boundaries where the free boundary is highlighted in red. The colour in (a)–(b) represents *q*(*x, t*) and 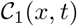 respectively. Discrete and continuum solutions for the average cell density, 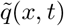, and the average chemical concentration, 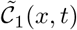 are compared in (c)–(d) respectively. The purple line in (c)–(d) shows the asymptotic behaviour of 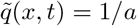 and 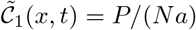 respectively.

Using Equation (97), *N* = *q*(*x,* 0)*L*(0) and 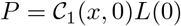. As the tissue relaxes, the cells elongate such that the length of individual cells approaches the cell resting length. Thus, *q*(*x, t*), 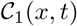 and *L*(*t*) become constant,

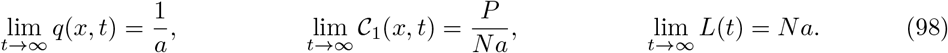

Figure 9(c)–(d) shows that the average cell density, 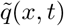, and the average chemical concentration, 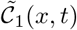, approach the limits stated in Equation (98) as *t* → ∞. The free boundary in Figure 9(a)–(b) demonstrates that *L*(*t*) approaches the limit stated in Equation (98) as *t* → ∞.

## E Case study 1: Rac–Rho pathway

To investigate the Rac–Rho pathway, we let 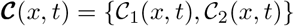 such that 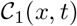 represents the concentration of RhoA and 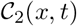 represents the concentration of Rac1. Figure 10 compares the discrete and continuum solutions relating to Figure 5.

**Figure 10:**
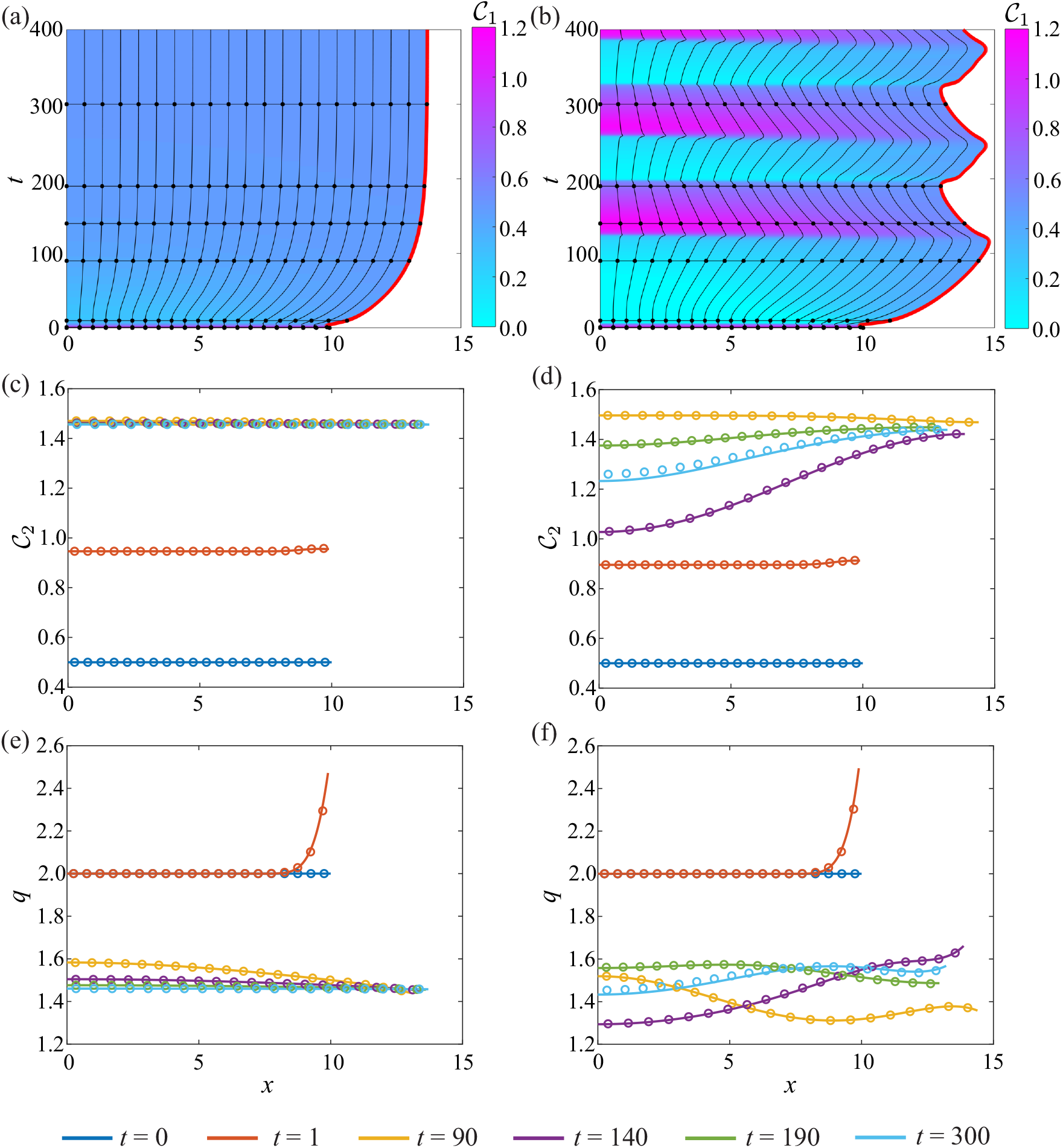
1D tissue dynamics for the interaction of RhoA and Rac1. (a),(c),(e) correspond to non-oscillatory system where 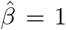 and (b),(d),(f) relate to an oscillatory system where 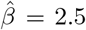. Characteristic diagrams in (a)–(b) illustrate the behaviour of 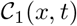 where the free boundary is highlighted in red. The black horizontal lines indicate times at which 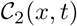 and *q*(*x, t*) snapshots are shown in (c)–(d) and (e)–(f) respectively. In (c)–(f), the discrete and continuum solutions are compared as the dots and solid line respectively for *t* = 0, 1, 90, 140, 190, 300. In both systems, 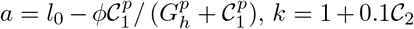, *D*_1_ = *D*_2_ = 1 and *η* = 1. Parameters: *b*_1_ = *b*_2_ = 1*, δ*_1_ = *δ*_2_ = 1*, n* = 3*, p* = 4, 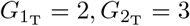, *l*_0_ = 1*, ϕ* = 0.65*, G*_*h*_ = 0.4.

## E.1 Single cell model

To investigate the influence of mechanobiological coupling on cellular dynamics, a discrete computational framework is used to model a single cell [12, 33]. We let **C**(*t*) = {*C*^(1)^(*t*)*, C*^(2)^(*t*)} where *C*^(1)^(*t*) represents RhoA and *C*^(2)^(*t*) represents Rac1. As only a single cell is of interest, **C**(*t*) is not indexed with a subscript (Figure 11). Mechanobiological coupling is introduced such that the cell resting length, *a* = *a* (**C**), and the cell stiffness, *k* = *k*(**C**), depend on the chemical family, **C**(*t*).

**Figure 11:**
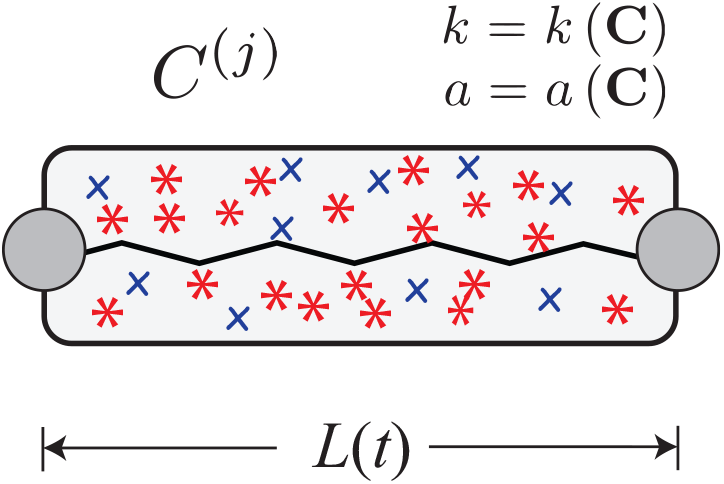
Schematic of a single cell with length *L*(*t*). The mechanical cell properties, *a* and *k*, depend on the family of chemical signals, **C**(*t*) = {*C*^(1)^(*t*)*, C*^(2)^(*t*)}.

We model the cell as an overdamped, mechanical spring [8, 40] such that **C**(*t*) tends to decrease as the cell expands, and tends to increase as the cell compresses. Thus, the governing equations are,

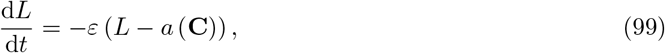

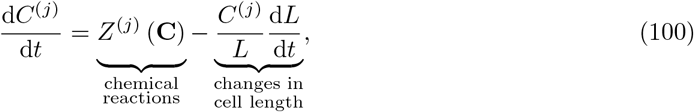

where *L*(*t*) is the cell length, *ε* = 2*k* (**C**) /*η* is the rate of contraction, *η* is the mobility coefficient and *Z*^(*j*)^ (**C**) governs the reactions between the chemical species within the cell [12]. For simplicity, the cell stiffness is chosen to be independent of **C**(*t*) such that *ε* is constant. The cell resting length is assumed to vary from a fixed value, *l*_0_ [12]. By including a Hill function with amplitude *ϕ*, switch location *G*_h_ and power *p*, we assume RhoA shortens the resting cell length [12],

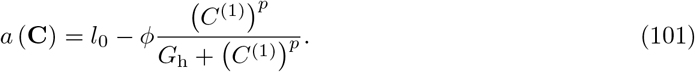

Equations (99) and (100) form a dynamical system. Phase planes are constructed to characterise the dependence of the system stability on the strength of the mechanobiological coupling. We used this analysis to inform our choice of model parameters in Section 3.2. The equilibrium points, 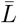 and 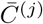, are determined by setting the time derivatives to zero such that 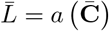 and 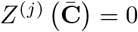.

To investigate a single cell containing only RhoA, we consider **C**(*t*) = {*C*^(1)^(*t*)} and let [12]

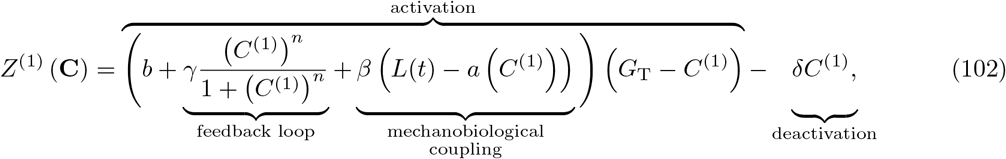

where *b* is the basal activation rate, *γ* is rate of feedback activation, *β* governs the strength of the mechanobiological coupling, *G*_T_ is the total amount of active and inactive RhoA, and *δ* is the deactivation rate [12, 33]. By substituting 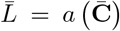 into 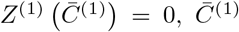 is numerically computed as the roots of,

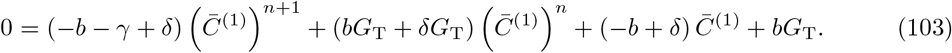

Equation (103) is independent of *β*. Thus, we vary *β* to investigate how the system stability depends on the strength of the mechanobiological coupling (Figure 12). Phase planes are constructed using `quiver` in MATLAB [67], and trajectories are computed using `ode15s` in MATLAB [43] (Figure 12).

**Figure 12:**
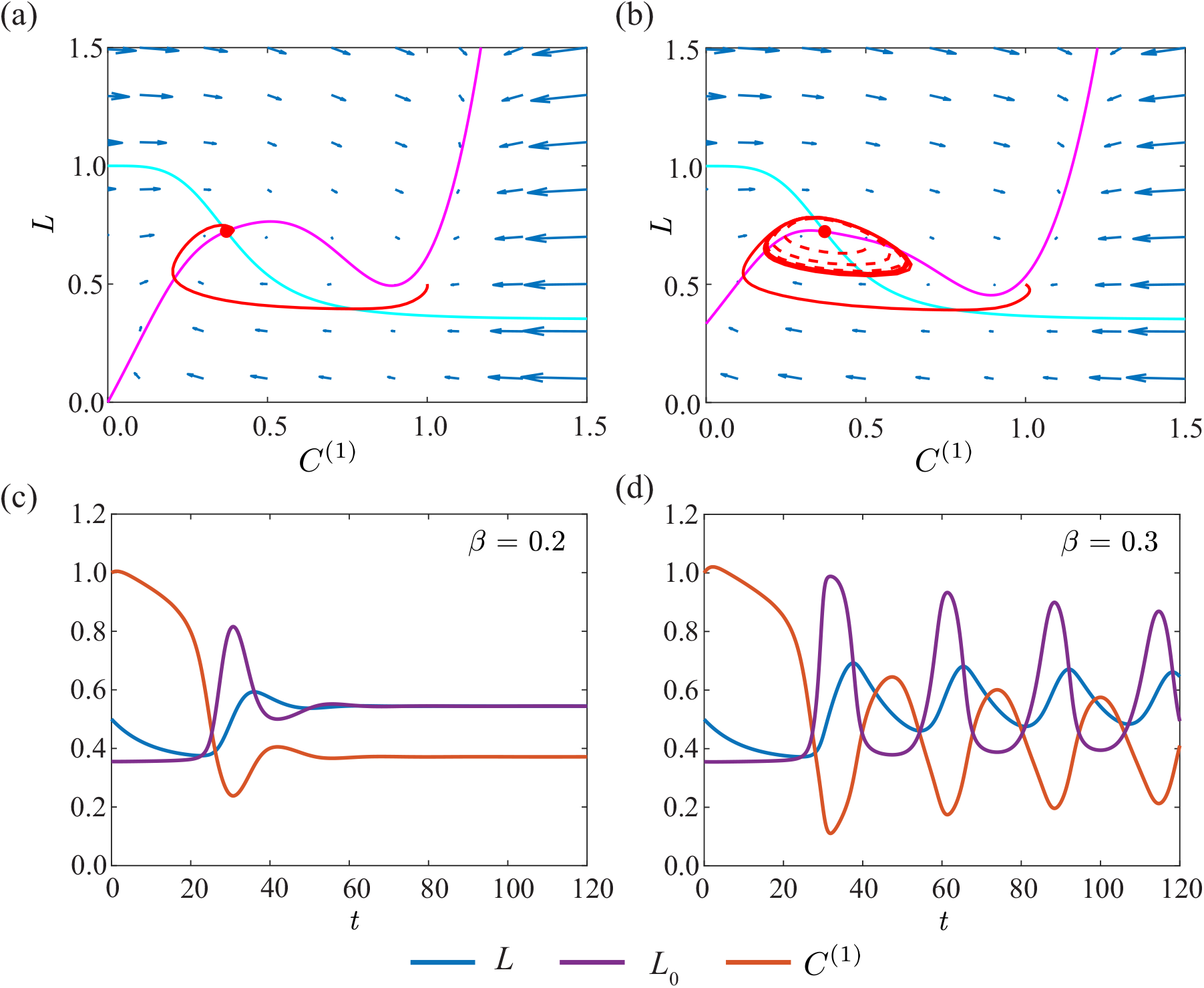
Dynamics of a single cell containing RhoA. (a),(c) corresponds to a non-oscillatory system where *β* = 0.2, and (b),(d) corresponds to an oscillatory system where *β* = 0.3. In (a)–(b), cyan represents the *L* nullcline and magenta represents the *C*^(1)^ nullcline. The trajectory for *C*^(1)^(0)*, L*(0) = (0.5, 1) is shown in red. In (b), the additional trajectory for *C*^(1)^(0)*, L*(0) = (0.5, 0.65) is drawn as a dashed red line to demonstrate that all solutions exhibit oscillatory behaviour for *β* = 0.3. Parameters are: *b* = 0.2*, γ* = 1.5*, n* = 4*, p* = 4*, G*_T_ = 2*, l*_0_ = 1*, ϕ* = 0.65*, G*_h_ = 0.4*, δ* = 1*, ε* = 0.1.

Figure 12(a),(c) demonstrates that the equilibrium point is stable when *β* = 0.2, and the cell exhibits non-oscillatory behaviour. By increasing the strength of the mechanobiological coupling to *β* = 0.3, a limit cycle arises, which leads to continuous oscillations in *L*(*t*) and *C*^(1)^(*t*) (Figure 12(b),(d)). Stability analysis reveals that the equilibrium point is unstable for *β* = 0.3. Thus, all solutions, regardless of the initial condition, exhibit oscillatory behaviour when *β* = 0.3 (Figure 12(b)).

To investigate how intracellular reactions between RhoA and Rac1 impact cell behaviour, we consider **C**(*t*) = {*C*^(1)^(*t*)*, C*^(2)^(*t*)} and let [12],

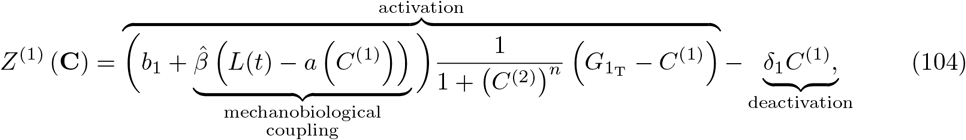

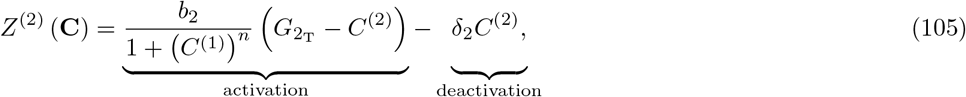

where *C*^(1)^(*t*) is the concentration of RhoA and *C*^(2)^(*t*) is the concentration of Rac1. Figure 13(a),(c) illustrates that when the mechanobiological coupling is weak, 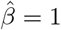, the equilibrium point is stable and the cell mechanically relaxes. By increasing the strength of the mechanobiological coupling to 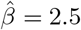, a limit cycle arises and the cell exhibits oscillatory behaviour (Figure 13(b),(d)).

**Figure 13:**
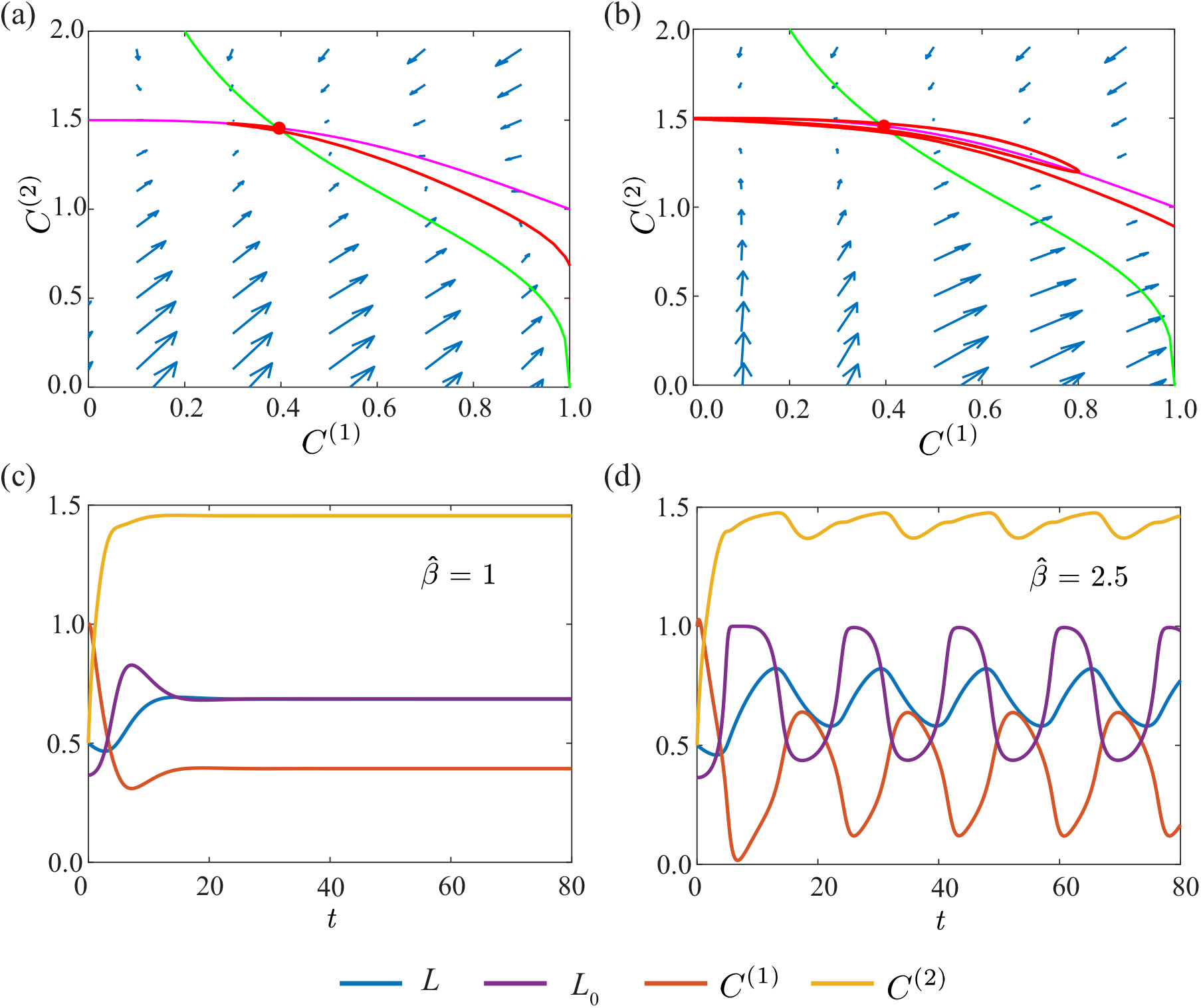
Reactions between RhoA and Rac1 in a single cell. (a),(c) corresponds to a non-oscillatory system where 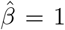, and (b),(d) corresponds to an oscillatory system where 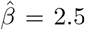. In (a)–(b), magenta represents the *C*^(1)^ nullcline and green represents the *C*^(2)^ nullcline. The trajectory for *L*(0)*, C*^(1)^(0)*, C*^(2)^(0) = (0.5, 1, 0.5) in shown in red. Parameters: *b*_1_ = *b*_2_ = 1*, δ*_1_ = *δ*_2_ = 1*, n* = 3*, p* = 4, 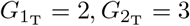, *l*_0_ = 1*, ϕ* = 0.65*, G*_h_ = 0.4*, ε* = 0.1.

## F Case study 2: Activator–inhibitor patterning

Section 3.3 considers an activator–inhibitor system with 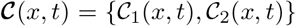, where 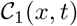 is the activator and 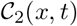 is the inhibitor. Figure 14 illustrates the long time behaviour of Figure 6(b),(d). Figure 14 demonstrates that two distinct activator peaks evolve as *t* → ∞ when *d > d*_*c*_. Thus, unlike [25, 26], we do not observe continuous peak splitting.

**Figure 14:**
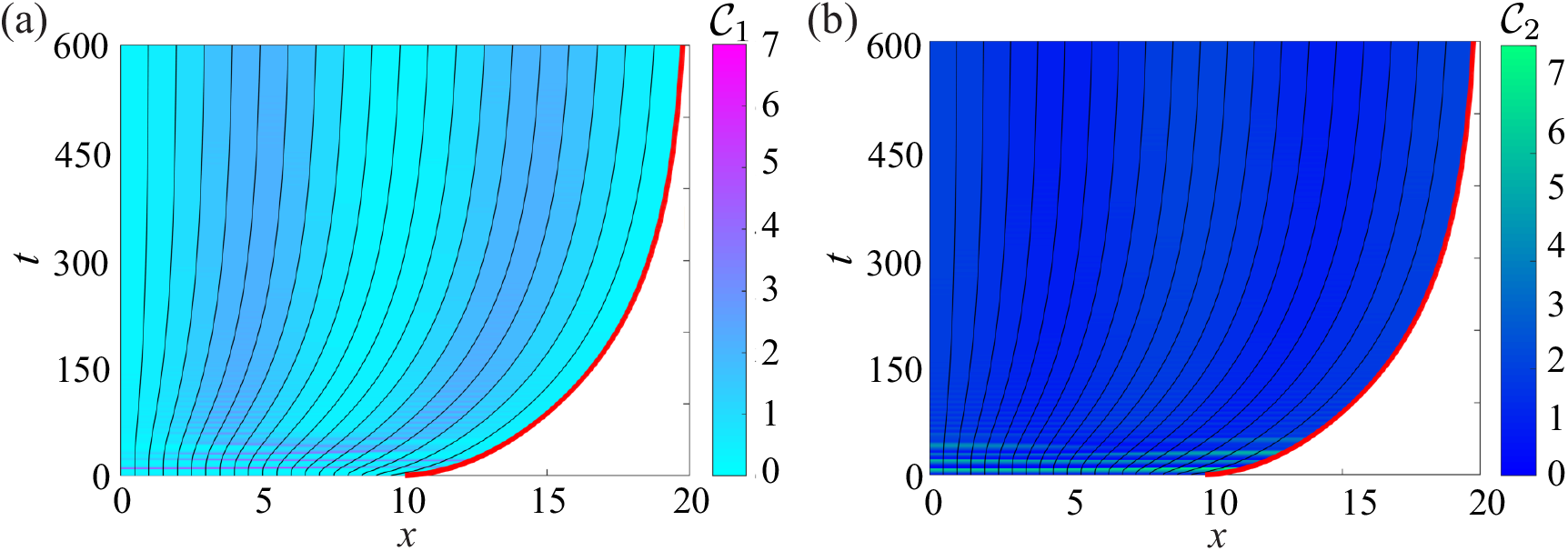
The evolution of spatial–temporal patterns in a homogeneous tissue with Schnakenberg dynamics, where *D*_1_ = 0.5 and *D*_2_ = 5 such that *d > d*_*c*_. (a) illustrates the behaviour of the activator, 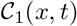, and (b) illustrates the behaviour of the inhibitor, 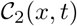. Parameters are as in Figure 6.

## F.1 Linear stability analysis and computation of critical diffusivity

In this section, we summarise the linear stability analysis used to determine the critical diffusivity value, *d*_*c*_. The classical Turing analysis for reaction–diffusion equations on fixed domains is outlined in Chapter 2 of [55]. As this paper considers reaction–diffusion equations on evolving cellular domains, our approach to the linear stability analysis is based on the classical Turing analysis.

Following [55], we non-dimensionalise the governing equations (Equations (26) and (30)). For the activator–inhibitor system stated in Section 3.3, the governing equations for a homogeneous tissue are:

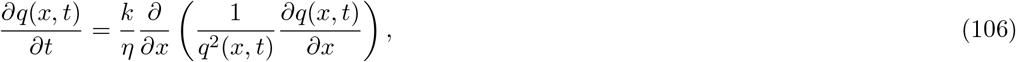

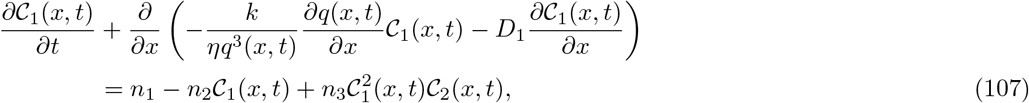

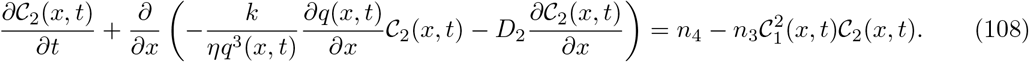

To non-dimensionalise Equations (106)–(108), we consider:

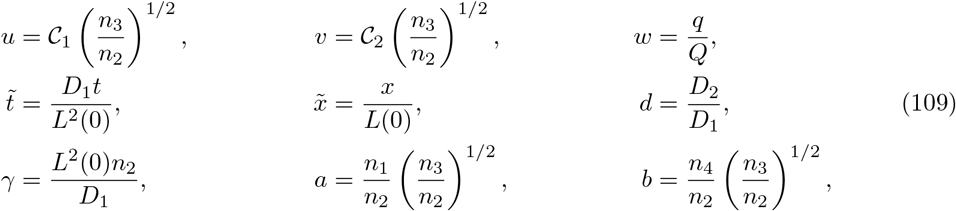

where *Q* has the same magnitude and units of *q*(*x, t*). Setting 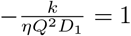 gives the non-dimensional governing equations as,

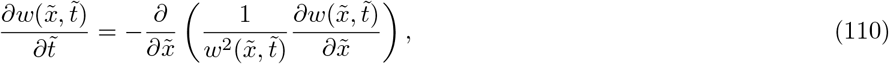

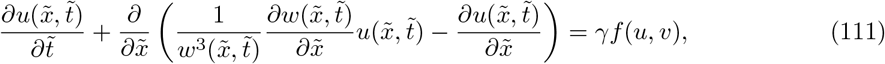

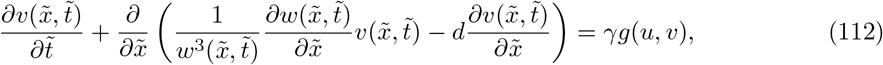

where

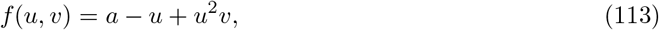

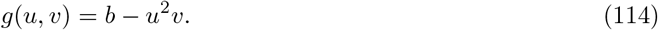

As setting the spatial derivatives in Equations (111)–(112) to zero yields Equation (2.13) of [55], the linear stability analysis is similar to the classical Turing analysis [55].

The classical Turing analysis considers the conditions where the homogeneous steady state, (*u, v*) = (*u*_0_*, v*_0_), is linearly stable in the absence of spatial variation [55]. Setting Equations (113)–(114) to zero gives the homogeneous steady state as *u*_0_ = *a* + *b* and *v*_0_ = *b/*(*a* + *b*)^2^. For Turing patterns on fixed domains, Equation (2.27) of [55] states that the critical diffusivity is the positive root of

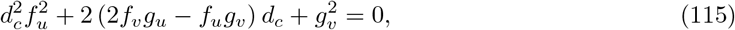

where the derivatives in Equation (115) are evaluated at (*u*_0_*, v*_0_). Using the quadratic formula, the critical diffusivity for Turing patterns on fixed domains is

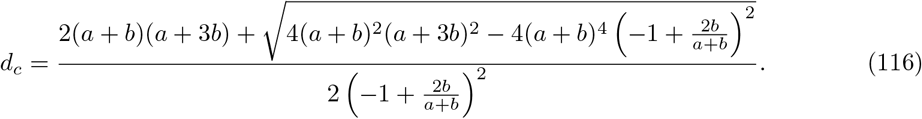

Guided by Equation (116), we explore relative diffusivity values above and below *d*_*c*_ to observe the formation of spatial–temporal patterns on an evolving cellular domain.

